# Mitochondrial DNA removal is essential for sperm development and activity

**DOI:** 10.1101/2024.05.07.592931

**Authors:** Zhe Chen, Fan Zhang, Annie Lee, Michaela Yamine, Zong-Heng Wang, Guofeng Zhang, Chris Combs, Hong Xu

**Affiliations:** National Heart, Lung, and Blood Institute, NIH; National Institute of Biomedical Imaging and Bioengineering, NIH

**Author notes:** Correspondence: Hong Xu.

## Abstract

During multicellular organisms’ reproduction, organelles, cytoplasmic materials, and mitochondrial DNA (mtDNA) are all derived from maternal lineage. Active mtDNA elimination during spermatogenesis has emerged as a conserved mechanism ensuring the uniparental mitochondrial inheritance in animals ^1 2 3 4^. However, given the existence of post-fertilization processes degrading sperm mitochondria ^5 6 7 8 9^, the physiological significance of sperm mtDNA removal is not clear. We report that mtDNA removal is indispensable for sperm development and activity. We uncover a novel mitochondrial exonuclease, ExoA (Exonuclease A) that is specially expressed in late spermatogenesis and exclusively required for mtDNA elimination. Loss of ExoA impairs mtDNA clearance in elongated spermatids and impedes the progression of individualization complexes that strip away cytoplasmic materials and organelles. Additionally, persistent mtDNA in mature sperm causes marked fragmentation of nuclear genome and complete sterility of *exoA* mutant male flies. All these defects can be suppressed by expressing a mitochondrially targeted bacterial exonuclease to ectopically remove mtDNA. Our work illustrates the developmental necessity of mtDNA clearance for the effective cytoplasm removal at the end of spermatid morphogenesis and to prevent potential nuclear-mito imbalance in mature sperms, in which the activity of nuclear genome is shutdown. Hence, the uniparental mitochondrial inheritance seems co-evolved with a key feature of sexual reproduction, the asymmetry between two gametes.

## Main

Mitochondrial genome is transmitted exclusively through maternal lineage in most sexually reproduced organisms. However, the underlying mechanisms of uniparental inheritance are not well-understood, and its physiological significance remains elusive. The maternal inheritance was once attributed to the simple dilution of sperm mitochondrial DNA (mtDNA) in the zygote, which contains an enormous amount of maternal mtDNA ^10,11^. However, increasing evidence suggests that mtDNA and mitochondria-derived structures are actively eliminated during spermatogenesis ^1 2 3 4^ or embryogenesis ^5 6 7 8 9^, respectively. Considering the high energy demand of the spermatogenesis process and sperm motility, it is intriguing why fathers proactively remove sperm mtDNA before fertilization. Understanding the evolutionary drive of mtDNA removal in spermatogenesis is of great interest. In *Drosophila* testis, abundant mitochondrial nucleoids are observed along the length of mitochondrial derivatives in elongating spermatids *^2^*, but the majority of them abruptly disappear in fully elongated spermatids. Few remaining nucleoids are stripped away by progressing actin cones during spermatid individualization, and eventually end up in waste bags. EndoG ^2^, a mitochondrial endonuclease, and Tamas ^3^, the mitochondrial DNA polymerase, are involved in the pre-individualization mtDNA removal. EndoG is a site-specific endonuclease, targeting both mtDNA ^12 13^ and nuclear DNA (nuDNA) ^14^. Notably, the polymerase activity of Tamas, not its exonuclease activity, is required for mtDNA removal ^3^. Hence, additional nucleases, particularly an exonuclease, are likely involved in further degrading EndoG-nicked mtDNA. Additionally, the pleiotropy of EndoG and Tamas poses challenges to understanding the physiological significance of mtDNA removal during spermatogenesis.

### A testis mitochondrial nucleoid protein, ExoA, is required for male fertility

We hypothesized that a protein specifically involved in mtDNA removal would likely show biased expression in testis and be associated with mitochondrial nucleoids. To this end, we surveyed the expression patterns of *Drosophila* homologs of previously identified mammalian nucleoid proteins ^15 16 17^ (Extended Data Fig. 1; Supplementary table 1). Among them, a candidate nucleoid protein encoded by the *CG12162* locus, known as Poldip2, was found to be highly enriched in testes (Extended Data Fig. 1). Poldip2 was initially identified as a human polymerase δ P50-interacting protein in a yeast two-hybrid screen ^18^ and has been proposed to be involved in nuclear genome replication and repair^19^. However, Poldip2 was exclusively localized to the mitochondrial matrix based on a GFP complementation assay (Fig. 1a) and concentrated on mitochondrial nucleoids (Fig. 1b). Therefore, Poldip2 is unlikely to interact with polymerase δ, a nuclear protein. To avoid future confusion, we renamed Poldip2 to ExoA due to its exonuclease activity, as described below.

**Fig. 1.**
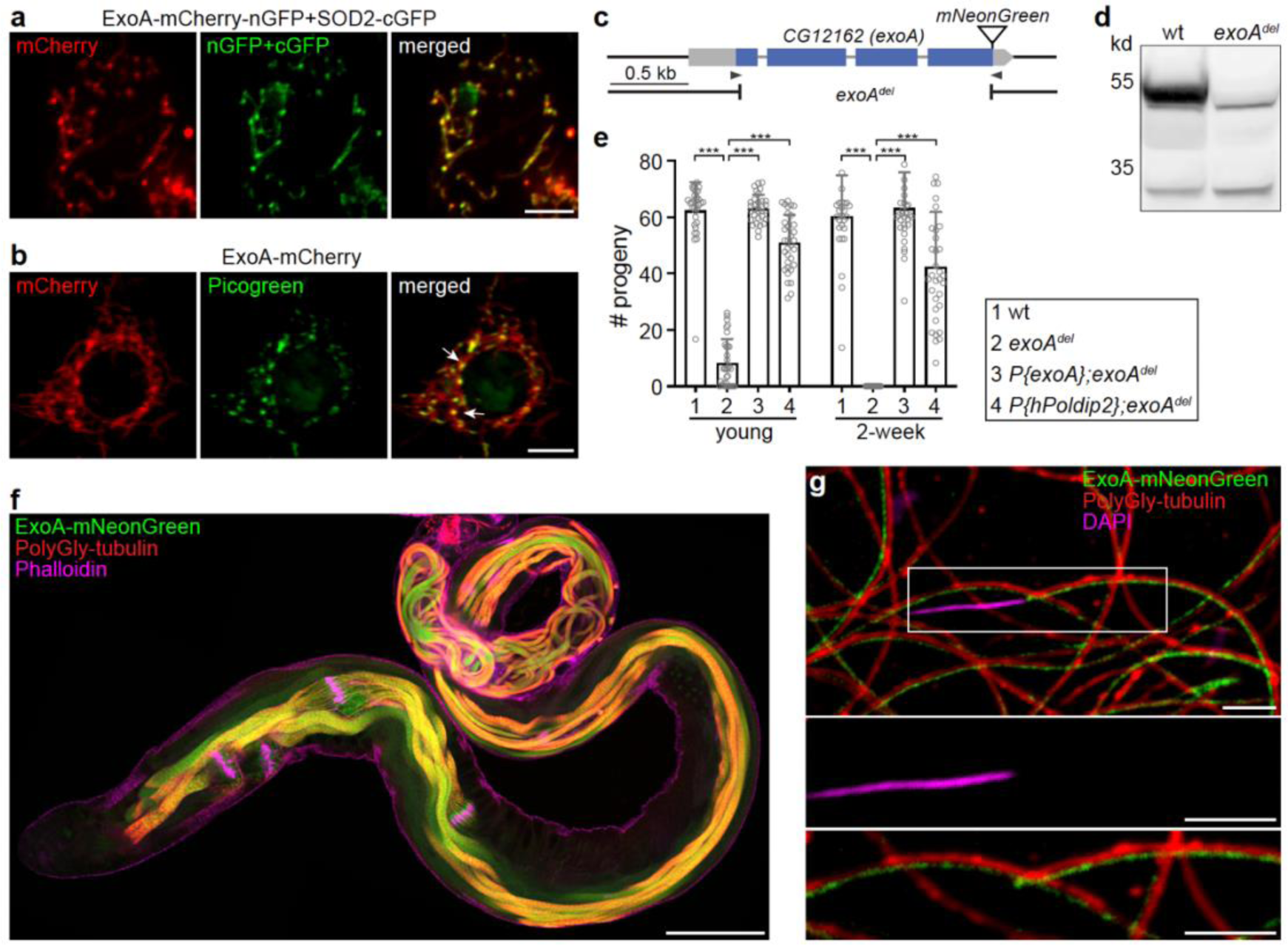
A mitochondrial nucleoid protein, ExoA is required for male fertility. **a**, A representative image of S2 cells co-expressing ExoA-mCherry-nGFP (red) and SOD2-cGFP. Two half-GFP molecules reconstitute into a functional whole GFP (green), demonstrating that ExoA is co-localized with SOD2 in mitochondrial matrix. Bar, 5 μm. **b**, ExoA concentrates on specific loci on mitochondria which stain positively for Picogreen (arrows), indicating ExoA is associated with mitochondrial nucleoids. Bar, 5 μm. **c**, Schematic representation of the *CG12162* genomic locus, illustrating *CG12162* transcripts, 5’- and 3’-UTR (grey bars), exons (blue bars), and the deleted region in *exoA^del^*. Arrows illustrate the target sites of guide RNAs used for generating the *exoA* deletion, and for knocking in *mNeonGreen*. **d**, Western blot confirms the deletion of the ExoA protein in *exoA^del^*flies. **e**, The *exoA^del^* flies are male semi-sterile. The number of progenies produced per day are shown. Compared with wild-type, male *exoA^del^* flies produce significantly fewer progeny at a young age and become completely sterile after two weeks. The semi-sterile phenotype can be rescued by expressing either ExoA or hPoldip2 protein in *exoA^del^* flies. Error bars represent standard deviation. *** P<0.0001. **f**, ExoA is highly expressed in fully elongated spermatids and all subsequent developmental stages within *Drosophila* testes. Polyglycylated tubulin marks fully elongated axonemal microtubules. Phalloidin stains actin cones and outlines the testis. Bar, 100 μm. **g**, Representative image shows that ExoA aligns alongside microtubules and is absent from the nuclear head region in spermatozoa, demonstrating that ExoA exclusively localizes in mitochondria in mature spermatozoa (enlarged view outlined). Green: ExoA-mNeon-Green; Red: Polyglycylated tubulin; Magenta: DAPI; Bar, 5 μm.

Using CRISPR/Cas9-mediated non-homologous end joining, we generate a deletion, *exoA^del^*, which removed most the coding region of *exoA* (Fig. 1c). No ExoA protein was detected in *exoA^del^* flies (Fig. 1d), indicating this deletion is a null allele. The *exoA^del^* flies were largely healthy, except that male flies were semi-sterile (Fig. 1e). Newly-eclosed male *exoA^del^* flies produced much less progeny compared to wild-type (wt) flies and became completely sterile after two weeks (Fig. 1e). The fertility of *exoA^del^* male flies was fully restored by a mini gene spanning the genomic region of *exoA* (Fig. 1e), demonstrating that impaired male fertility is caused by the loss of ExoA. Additionally, a transgene containing the cDNA of human homolog of *exoA*, *hPoldip2*, flanked by 5’- and 3’-UTRs of *exoA*, largely restored the fertility of *exoA^del^* male flies, indicating a conserved function of ExoA in maintaining male fertility in metazoans (Fig. 1e).

### Persistent mtDNA impairs male fertility

To elucidate the underlying cause of impaired fertility of *exoA^del^* male flies, we examined the developmental expression pattern of ExoA in the testes. We generated an ExoA-mNeonGreen (ExoA-mNG) reporter by inserting the *mNeonGreen* cDNA into the endogenous locus of *exoA* (Fig. 1c). ExoA-mNG expression was hardly detected in early spermatogenesis stages but exhibited an abrupt increase in fully elongated spermatids, which were marked with polyglycylated tubulin (Fig. 1f). Its expression persisted through all subsequent developemetal stages, including mature spermatozoa desposited in seminal vesicles (Fig. 1g).

The onset of ExoA expression coincided with mtDNA removal in elongated spermatids ^2^. Additionally. ExoA was found to be enriched on mitochondrial nucleoids (Fig. 1b). These observations intrigued us to test whether ExoA is involved in mtDNA removal. We stained isolated spermatid bundles with DAPI, a fluorescent DNA dye, to visualize mtDNA. In elongating spermatids, mitochondrial nucleoids were evenly distributed in both wt and *exoA^del^* sperm tails (Fig. 2a). The total number of nucleoids in early- and mid-elongating spermatids was comparable between wt and *exoA^del^* (Fig. 2d and Extended Data Fig. 2a), although some nucleoids were notably larger in *exoA^del^* spermatids (Extended Data Fig. 2c), suggesting a potential defect in nucleoid organization. In fully elongated spermatids, most mitochondrial nucleoids disappeared in wt (Fig. 2b, d, e and Extended Data Fig. 2a, b), indicating active mtDNA removal at this stage. In contrast, a significant amount of mtDNA remained in *exoA^del^* spermatids (Fig. 2b, d, e and Extended Data Fig. 2a, b) and persisted after individualization (Fig. 2c). As intact seminal vesicles are impermeable to DAPI, we imaged an endogenously tagged TFAM reporter, TFAM-mNeonGreen ^20^, which marks mitochondrial nucleoids (Extended Data Fig. 2d, e), to assess mtDNA in mature sperm. Many TFAM puncta were detected inside *exoA^del^* seminal vesicles, while no nucleoids were found in wt seminal vesicles (Fig. 2f).

**Fig. 2.**
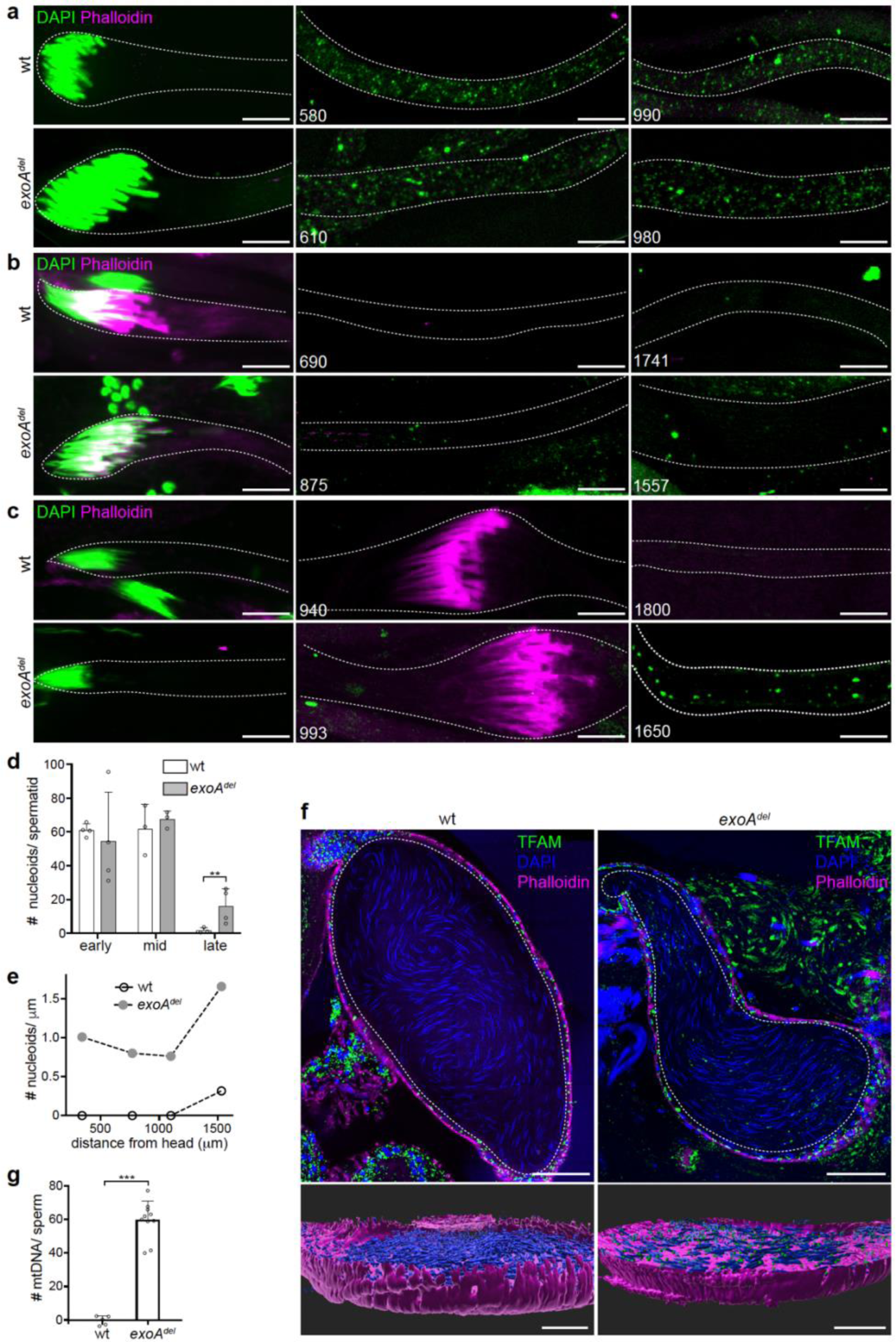
mtDNA persist in late spermatogenesis stages of *exoA^del^*flies. **a-c**, Representative images of elongating (**a**), fully elongated (**b**) and individualization (**c**) spermatid bundles isolated from *w^1118^* (wt) and *exoA^del^* flies, and stained for DNA (DAPI, green) and actin cones (Phalloidin, magenta). Note that DAPI stains both nuclear DNA (nuDNA) and mitochondrial DNA (mtDNA) in isolated spermatid bundles. Dashed lines outline bundles. Numbers indicate the distance (μm) from the nuclear head. Bar, 10 μm. **d**, Quantification of the total mitochondrial nucleoid numbers per spermatid at early-elongating (early), mid-elongating (mid) and fully elongated (late) stages. Error bars represent standard deviation. ** P<0.001. **e**, Representative density of mitochondrial nucleoids (total numbers per μm) along the length of fully elongated spermatid bundles for *w^1118^* (wt) and *exoA^del^* flies, respectively. **f**, Representative images show numerous mitochondrial nucleoids labeled by TFAM-mNeonGreen in *exoA^del^,* but not wt mature sperms in seminal vesicle (dashed lines). 3D rendering of seminal vesicle. Phalloidin stains actin (magenta); DAPI stains needle-shaped nuDNA (blue). Bar, 50 μm. **g**, Droplet digital PCR (ddPCR) quantification of the average number of paternal mtDNA molecules per sperm in the female spermatheca. Crosses were performed between female *w^1118^* (*mt:ND2^del1^*) flies and male *w^1118^* (*mt:wt*) or *exoA^del^* (*mt:wt*) flies. Error bars represent standard deviation. *** P<0.0001. See also Extended Data Fig. 2g, g’.

We further quantified the copy number of mtDNA molecules in mature sperms. Either *w^1118^* or *exoA^del^* male flies carrying wild-type mtDNA (*mt:wt*) were mated with *w^1118^* female flies carrying homoplasmic *mt:ND2^del1^*, which contains a 9-base pair deletion on mtDNA-encoded *ND2* locus. Subsequently, we dissected the sperm storage organ, spermatheca, from copulated female flies and subjected them to droplet digital PCR analysis, using primers specifically targeting paternal mtDNA (*mt:wt*) (Extended Data Fig. 2f, f’). In the cross using *w^1118^* male flies, no paternal mtDNA was detected (Extended Data Fig. 2g, g’), supporting the notion that mature sperm are devoid of mtDNA in *Drosophila* ^2 3^. However, when fathers were *exoA^del^* flies, a significant amount of paternal mtDNA was detected (Extended Data Fig. 2g, g’). On average, each mature sperm of *exoA^del^* had 59 copies of mtDNA (Fig. 2g). Paternal mtDNA was detected in embryos 30 minutes after egg laying but disappeared 6 hours later (Extended Data Fig. 2h, h’), consistent with the notion that sperm mitochondria are destroyed during early embryogenesis ^8^. Altogether, these observations demonstrate that ExoA is essential for mtDNA removal in late spermatogenesis.

We next addressed whether the persistence of mtDNA in mature sperm is the cause of impaired fertility in *exoA^del^* male flies. Given that EndoG nicks mtDNA in developing spermatids ^2^, we reasoned that ectopically introducing an exonuclease into mitochondria would degrade mtDNA nicked by EndoG in *exoA^del^* spermatids. We replaced the coding region of *exoA* with a fusion gene consisted of a mitochondrial targeting sequence and the cDNA encoding *E.coli* Exonuclease III (*mitoExoIII*), using CRISPR/Cas9 mediated recombination (Fig. 3a). To prevent potential leaky expression of mitoExoIII, we placed the SV40 transcription-terminating sequence ^21^, flanked by two Flippase (FLP) recombination target (FRT) sites, in front of *mitoExoIII*. In the presence of FLP activated by a *Bam-gal4*, FRT-mediated recombination excises the termination sequence, allowing the expression of mitoExoIII under the control of *exoA* promoter exclusively in germline. Expression of mitoExoIII in the heteroallelic combination of the *exoA* null background reduced the abundance of remaining mitochondrial nucleoids in mature sperm (Fig. 3b), and importantly, restored male fertility in both young and old *exoA* null flies (Fig. 3c), indicating that the persistence of mtDNA in late spermatogenesis impairs male fertility.

**Fig. 3.**
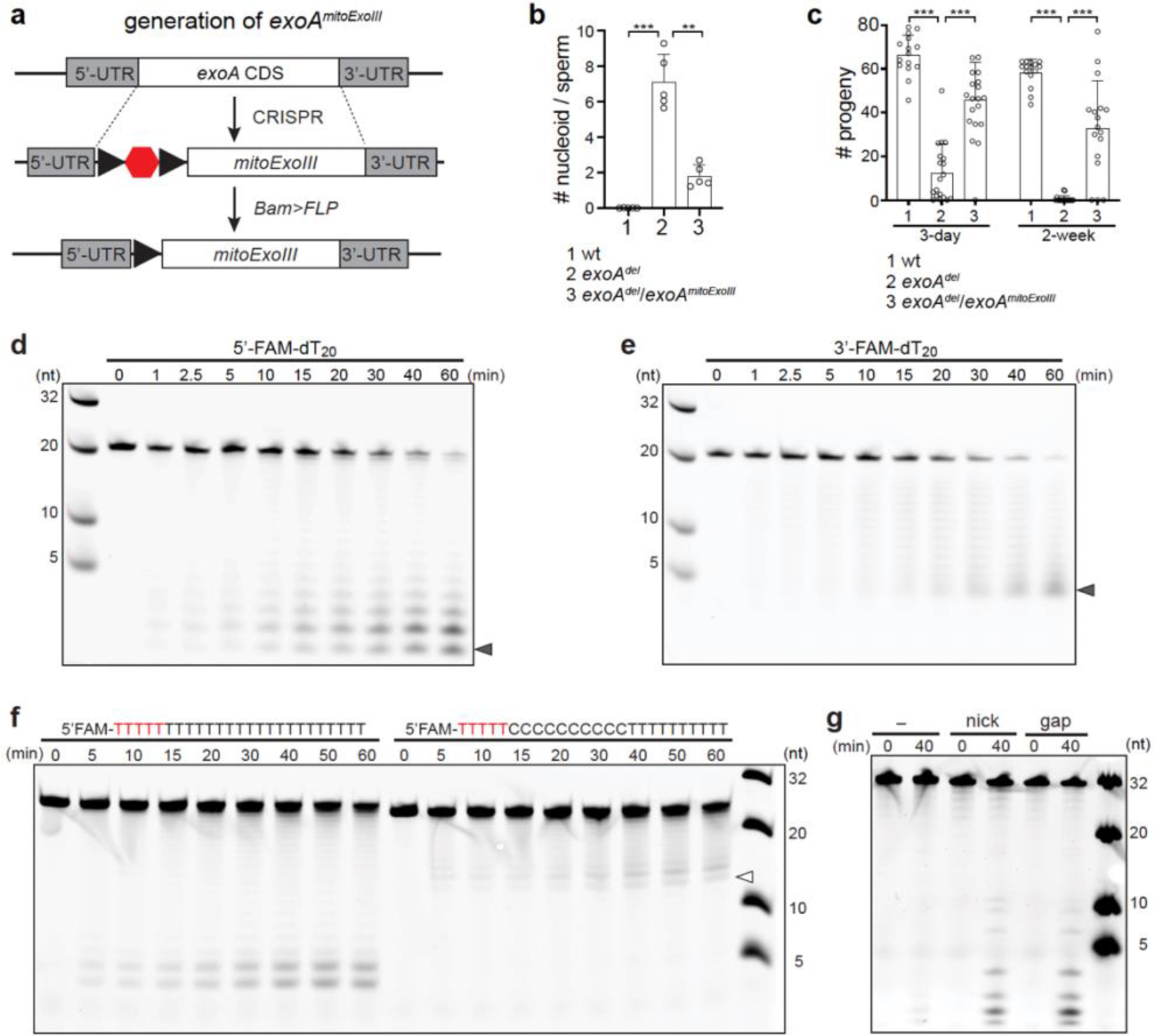
ExoA is a mitochondrial DNA exonuclease. **a**, Genetic scheme of replacing *exoA* coding sequence (CDS) with a mitochondrially targeted *E. coli* Exonuclease III (mitoExoIII) in developing spermatids (*exoA^mitoExoIII^*). The SV40 transcription termination sequence (hexagon) flanked by two FRT sites (arrowheads) allows conditional expression of mitoExoIII induced by Flippase (FLP). **b**, Expression of mitoExoIII in spermatids reduced mitochondrial nucleoid numbers in mature sperms of 3-day-old *exoA^del^* flies. 1. wt: *UAS-FLP*/+; *Bam-gal4*, *exoA ^del^ /+.* 2. *exoA ^del^*: +/+; *Bam-gal4*, *exoA ^del^*/*exoA^mitoExoIII^*; 3. *exoA ^del^*/*exoA^mitoExoIII^*: *UAS-FLP*/+; *Bam-gal4*, *exoA ^del^*/ *exoA^mitoExoIII^*. Error bars represent standard deviation. ** P<0.001; *** P<0.0001. **c**, Expression of mitoExoIII in spermatids rescued the fertility of both young and 2-week-old male *exoA^del^* flies. The number of progenies per day are shown. 1. wt: *UAS-FLP*/+; *Bam-gal4*, *exoA ^del^ /+.* 2. *exoA ^del^*: +/+; *Bam-gal4*, *exoA ^del^*/*exoA^mitoExoIII^*; 3. *exoA ^del^*/*exoA^mitoExoIII^*: *UAS-FLP*/+; *Bam-gal4*, *exoA ^del^*/ *exoA^mitoExoIII^*. Error bars represent standard deviation. *** P<0.0001. **d**, ExoA exhibits 3’-5’ exonuclease activity. A 5’-6-FAM labeled 20-nt poly(dT) (100 nM) was incubated with the ExoA protein (200 nM) at 37 °C and analyzed at indicated time points. A ladder-like pattern of oligonucleotides ranging from monomer (arrowhead) to 19-mer was generated. **e**, ExoA displays 5’-3’ exonuclease activity. A 3’-6-FAM labeled 20-nt poly(dT) (100 nM) was incubated with the ExoA protein (200 nM) at 37 °C and analyzed at indicated time points. The resulting products showed a ladder pattern ranging from 3-mer (arrowhead) to 19-mer. **f**, ExoA degrades dC less efficiently. The 5’-6-FAM labeled 25-nt poly(dT) or 25-nt poly(dTdC) (100 nM) was incubated with the ExoA protein (400 nM) at 37 °C and analyzed at indicated time points. The 5’-end five nucleotides were protected from degradation by incorporting the internucleotide phosphorothioate bonds (red). Note the smallest degradation products is 15-mer (arrowhead) in the reaction of poly(dTdC), suggesting the stretch of dC inhibits the progression of the exonuclease. **g**, ExoA degrades double-stranded (dsDNA) with breaks. Three types of dsDNA, including blunt-ended dsDNA (-), dsDNA with a single nick (nick), and dsDNA with a single gap (gap), were generated using a 5’-6-FAM-labeled, 32-nt long oligonucleotide. The dsDNA substrates (100 nM) were incubated with the ExoA protein (1200 nM) at 37 °C and analyzed after 40 min. The molecular markers in this figure are an equal molar mixture of 5’-6-FAM labeled 32-nt, 20-nt, 10-nt and 5-nt oligonucleotides and were loaded at a concentration of 50 nM for each.

### ExoA is a mitochondrial exonuclease

A previous study showed that Poldip2 diminished the signal of DNA probes ^19^, although this phenomenon has not been further investigated. Herein, we found that a foreign exonuclease can functionally replace ExoA in *Drosophila*. These observations prompted us to explore whether ExoA might have nuclease activity. We expressed and purified the full-length recombinant ExoA protein from *E.coli* to a purity above 96% (Extended Data Fig. 3a). Incubation of 5’-6-FAM labeled 20-nt poly(dT) (Fig. 3d) or poly(dA) (Extended Data Fig. 3b) with recombinant ExoA resulted in a ladder-like pattern of oligonucleotides ranging from monomer to 19-mer. ExoA also degraded 3’-6-FAM labeled 20-nt poly(dT) (Fig. 3e) or poly(dA) (Extended Data Fig. 3c) in the 5’-3’ direction. In a reaction using a single-stranded DNA (ssDNA) substrate consisting of mixed nucleotides (Extended Data Fig. 3d), different intensities of degradation intermediates were observed, suggesting a potential nucleotide preference of ExoA. Indeed, ExoA showed minimal degradation on a 20-nt poly(dC) (Extended Data Fig. 3e-h), and a stretch of tandem dC hindered the progression of ExoA on a ssDNA substrate (Fig. 3f). Furthermore, we examined ExoA’s exonuclease activity on double-stranded DNA (dsDNA) substrates with various configurations. While the intact dsDNA substrate exhibited minimal degradation by ExoA, dsDNA with either a nick or a gap was susceptible to degradation (Fig. 3g). Collectively, these results demonstrate that ExoA is a DNA exonuclease, which can degrade both ssDNA and dsDNA with breaks, and prefers dA/dT over dG/dC. We hence renamed the *CG12162* locus, previously known as *Drosophila Poldip2* to *exonuclease A* (*exoA*).

### Persistent mtDNA impedes individualization

We have established that ExoA is a mitochondrial exonuclease highly enriched in the testis, specifically degrading mtDNA in elongated spermatids. We next explored how the presence of mtDNA impairs male fertility in *exoA^del^* flies. The testis of *exoA^del^* flies did not exhibit obvious morphological defects (Extended Data Fig. 4a, f). Consistent with ExoA’s spatial pattern, early stages of spermatogenesis seemed unaffected in *exoA^del^* testes. While the formation of individualization complexes (ICs), traveling ICs and waste bags also appeared normal in *exoA^del^* testes (Extended Data Fig. 4, b-d, g-i), the coiling region was notably enlarged (Fig. 4a and Extended Data Fig. 4k), accumulating many needle-shaped nuclei, some of which remained bundled together (Fig. 4a), suggesting a potential individualization defect. The seminal vesicles of both 3-day and 2-week-old *exoA^del^* flies were notably smaller and contained fewer mature sperm compared to wt (Fig. 4b and Extended Data Fig. 4e, j). In transmission electron microscopic analysis of the individualized cyst of *exoA^del^* flies, some spermatids were still enveloped in a continues membrane structure, aside from the remaining spermatids that were properly individualized (Fig. 4c, red arrowhead). Additionally, individualized spermatids showing incomplete membrane contour were frequently observed (Fig. 4c, red arrows). Both phenotypes suggest a potential defect in sperm individualization.

**Fig. 4.**
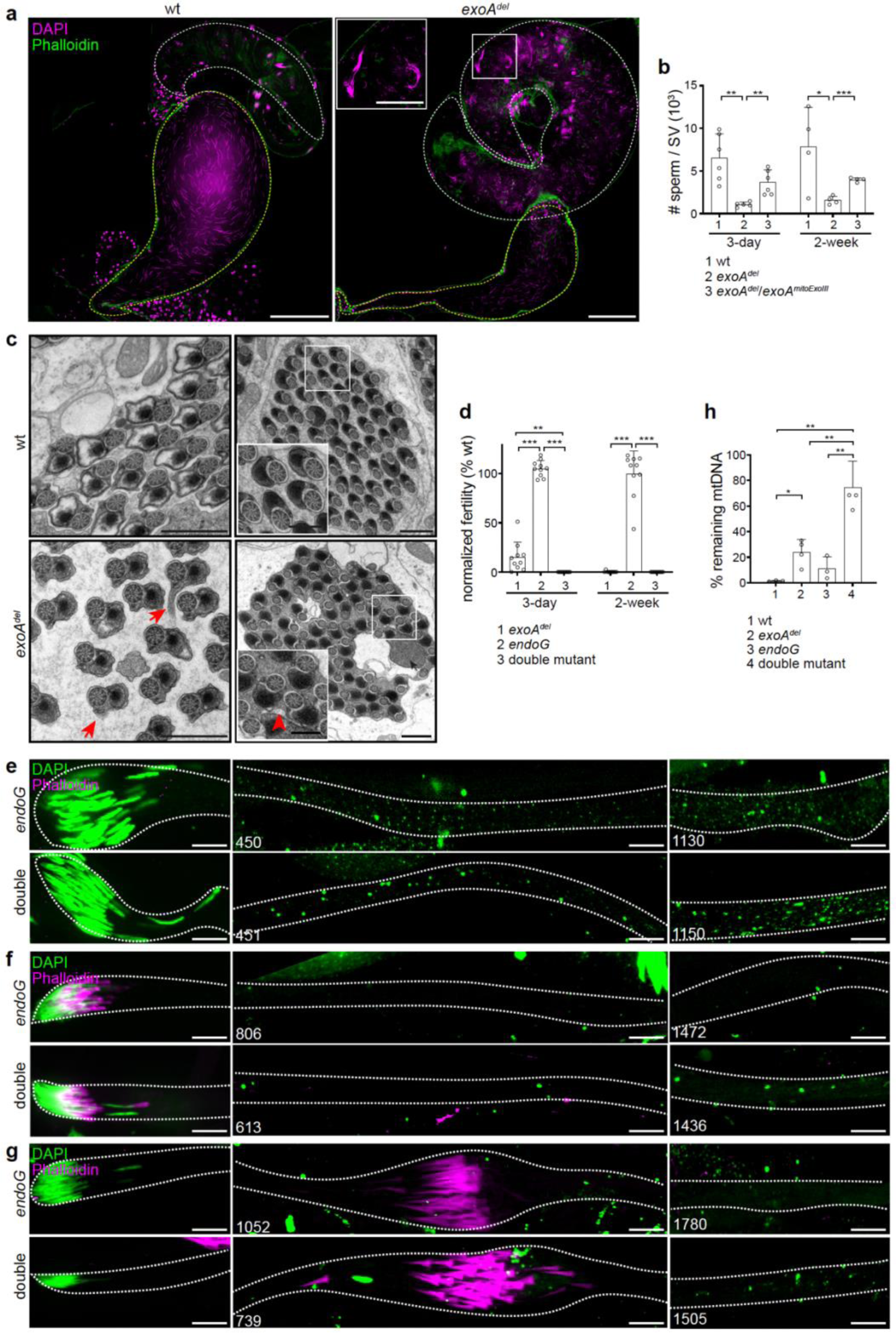
Persistent mtDNA impedes spermatid individualization. **a**, Representative images showing the coiling region (white dashed line) and seminal vesicle (yellow dashed line) of *w^1118^* (wt) and *exoA^del^* testes. Bar, 100 μm. Inset, the coiling region of *exoA^del^* testes accumulate many needle-shaped nuclei stained with DAPI (magenta), some of which remained bundled together. Phalloidin: green. Bar, 50 μm. **b**, Fewer mature sperm are present in the seminal vesicles of both young and 2-week-old *exoA^del^* flies compared with wild-type control, a deficiency that can be rescued by expressing mitoExoIII in spermatids. The number of mature sperm nuclei was quantified in each seminal vesicle of indicated genotypes. 1. wt: *UAS-FLP*/+; *Bam-gal4*, *exoA ^del^ /+.* 2. *exoA ^del^*: +/+; *Bam-gal4*, *exoA ^del^*/*exoA^mitoExoIII^*; 3. *exoA ^del^*/*exoA^mitoExoIII^*: *UAS-FLP*/+; *Bam-gal4*, *exoA ^del^*/ *exoA^mitoExoIII^*. Error bars represent standard deviation. *** P<0.0001; ** P<0.001; * P<0.05. **c**, Representative transmission electron microscopy (TEM) images of cross sections of *Drosophila* testes from 3-day-old flies showing individualized spermatid cysts. In *exoA^del^* testes, red arrows denote spermatids with incomplete membrane contour; red arrowheads denote two connected spermatids; black arrows denote abnormal mitochondrial derivate structures. Bar, 1 μm. Bar in insets, 500 nm. **d**, The double mutant was completely sterile, in contrast to the normal fertility of *endoG* mutant. The fertility of male *exoA^del^, endoG* (*endoG^MB07150/KO^*) and double mutant (*endoG^MB07150/KO^; exoA^del^*) was normalized to that of *w^1118^* male flies and plotted. Error bars represent standard deviation. *** P<0.0001; ** P<0.001. **e-g**, Representative images showing the isolated spermatid bundles of the elongating (**e**), fully elongated (**f**) and individualization (**g**) stages stained for DNA (DAPI) and actin cones (Phalloidin) in *endoG* (*endoG^MB07150/KO^*) and double mutant (*endoG^MB07150/KO^*; *exoA^del^*). DAPI stains both nuDNA and mtDNA in isolated spermatid bundles. Note the disorganized actin cone structures in the double mutant. Dashed lines mark bundle boundary. Numbers indicate the distance (μm) from the anterior tip of the spermatid. Bar, 10 μm. **h**, Quantification of remaining mitochondrial nucleoids in fully elongated spermatids of *w^1118^*(wt), *exoA^del^, endoG* (*endoG^MB07150/KO^*) and double mutant (*endoG^MB07150/KO^*; *exoA^del^*) flies. Total mitochondrial nucleoids measured in volumes per spermatid in fully elongated stage were normalized to that of the elongating stage in each genotype. Error bars represent standard deviation. ** P<0.01; * P<0.05.

If persistent mtDNA impedes ICs progression, one would expect that a greater amount of remaining mtDNA would lead to a stronger individualization defect. ExoA, a mitochondrial exonuclease, might work in synergy with EndoG, a mitochondrial endonuclease, to rapidly eliminate mtDNA in elongated spermatids ^2^. Hence, we attempted to examine the spermatids individualization in a background lacking both *exoA* and *endoG*. We deleted the entire coding region of *endoG* using CRISPR technology and combined *exoA^del^*with a trans-heterozygous combination consisting of *endoG^KO^ and EndoG^MB07150^*. The double mutant (*endoG^MB07150/KO^*; *exoA^del^*) was completely sterile, contrasting with the normal fertility of trans *endoG* (*endoG^MB07150/KO^*) (Fig. 4d). Individual nucleoids were further enlarged in size on average (Fig. 4e-g and Extended Data Fig. 4l), indicating more severe defects in nucleoid organization or morphology. In elongated spermatids, approximately 74.5% of mitochondrial nucleoids persisted in double mutant, whereas 11.3% and 23.8% remained in the *endoG* mutant and *exoA^del^*, respectively (Fig. 4h). Importantly, the travelling ICs were disorganized, and abundant mtDNA was detected at both basal and distal regions of cystic bulges in the double mutant (Fig. 4g). The observation that more remaining mtDNA caused more severe individualization defects further substantiates that mtDNA removal is necessary for individualization, allowing the rapid and smooth progress of travelling ICs.

### Persistent mtDNA damages nuclear genome

Having demonstrated that mtDNA removal promotes sperm individualization, which explains the reduced fertility of young *exoA^del^* male flies, we next asked why older *exoA^del^* male flies were completely infertile. Serendipitously, we found that nuclear DNA (nuDNA) was markedly fragmented in mature sperm of 2-week-old *exoA^del^* flies, but not in young *exoA^del^* flies or wild-type flies at either age (Fig. 5a, b). Importantly, the ectopic expression of mitoExoIII, which can degrade mtDNA, suppressed the nuDNA fragmentation in mature sperms of old *exoA^del^* flies (Fig. 5b), indicating that the persistent mtDNA triggers nuDNA damage.

**Fig. 5.**
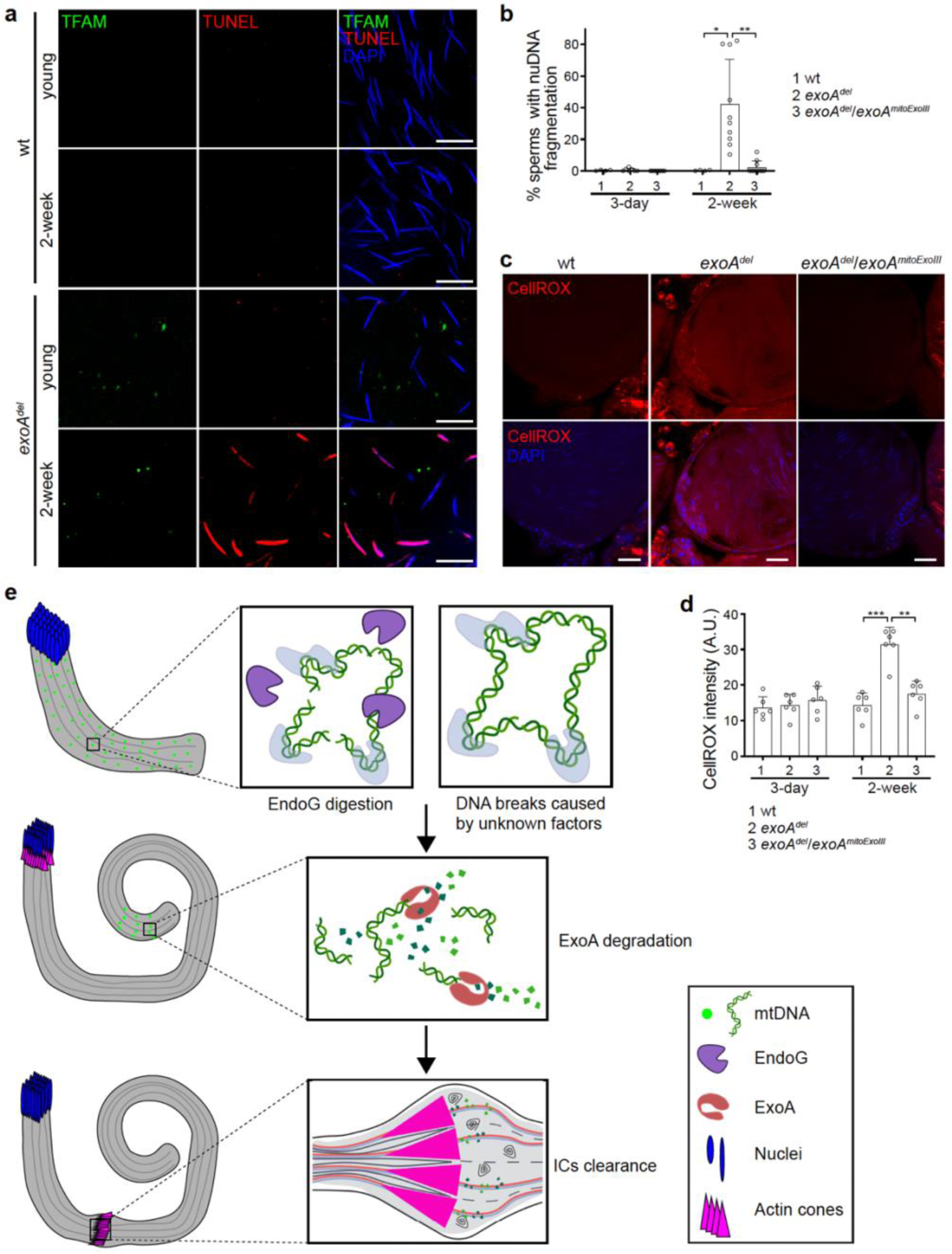
Persistent mtDNA in mature sperm damages nuclear genome. **a**, Representative images of TUNEL assay showing nuDNA breaks/fragmentation in 2-week-old *exoA^del^* mature sperm. TFAM-mNeonGreen labels mitochondrial nucleoids (green), and DAPI (blue) stains nuDNA of mature sperm in seminal vesicles. Red: TUNEL signal. Bar, 10 μm. **b**, Expression of mitoExoIII in spermatids reduces nuDNA breaks/fragmentation in *exoA^del^* mature sperm. Quantification was performed by normalizing the nuDNA breaks/fragmentation stained with TUNEL assay to the total sperm numbers stained by DAPI in a seminal vesicle. 1. wt: *UAS-FLP*/+; *Bam-gal4*, *exoA ^del^ /+.* 2. *exoA ^del^*: +/+; *Bam-gal4*, *exoA ^del^*/*exoA^mitoExoIII^*; 3. *exoA ^del^*/*exoA^mitoExoIII^*: *UAS-FLP*/+; *Bam-gal4*, *exoA ^del^*/ *exoA^mitoExoIII^*. Error bars represent standard deviation. ** P<0.001; * P<0.05. **c**, Representative images of CellROX Deep Red staining of 2-week-old fly sperms in the seminal vesicles of wt, *exoA^del^* and *exoA^del^* /*exoA^mitoExoIII^*flies. Red: CellROX Deep Red; Blue: nuDNA stained by DAPI. Bar, 20 μm. **d**, Quantification of CellROX intensity, as the measure for ROS level in both young and 2-week-old flies with indicated genotypes. 1. wt: *UAS-FLP*/+; *Bam-gal4*, *exoA ^del^ /+.* 2. *exoA ^del^*: +/+; *Bam-gal4*, *exoA ^del^*/*exoA^mitoExoIII^*; 3. *exoA ^del^*/*exoA^mitoExoIII^*: *UAS-FLP*/+; *Bam-gal4*, *exoA ^del^*/ *exoA^mitoExoIII^*. A.U., arbitrary unit. Error bars represent standard deviation. *** P<0.0001; ** P<0.001. **e**, Proposed model of pre-fertilization mtDNA removal. In elongating spermatids, mitochondria undergo dramatic structural changes, potentially sensitizing mitochondrial nucleoids. This may trigger the EndoG dependent mtDNA nicking or DNA breaks through other unknown mechanisms, initiating the clearance of mtDNA. In the final stage of spermatid elongation, the abrupt expression of ExoA leads to the complete degradation of mtDNA. During the individualization, individualization complexes (ICs) progress down the spermatids, gather any remaining oligonucleotides, and ultimately discard them in waste bags. Consequently, mature sperm are devoid of mtDNA. Created with BioRender.com.

During *Drosophila* spermatogenesis, nuDNA breaks occur to facilitate the histone-to-protamine transition from the post-meiotic stage onwards. These breaks are repaired in late elongation and sperm individualization stages. The timing of appearance and disappearance of nuDNA breaks were normal in *exoA^del^* flies (Extended Data Fig. 5a), suggesting that the nuDNA fragmentation is not caused by defects in chromatin remodeling. We stained the testes with tetramethylrhodamine methyl ester (TMRM) and MitoTracker Green, fluorescent dyes sensitive to mitochondrial membrane potential and mitochondrial mass, respectively. The ratio of TMRM intensity to that of Mitotracker Green, an indication of mitochondrial membrane potential, was decreased in mature sperm of young *exoA^del^* flies compared to the wt flies (Extended Data Fig. 5b). This reduced membrane potential was more pronounced in 2-week-old *exoA^del^* flies (Extended Data Fig. 5c). Correspondingly, reactive oxygen species (ROS), visualized with CellROX staining, were markedly increased in mature sperm of 2-week-old *exoA^del^* flies (Fig. 5c, d). It is likely that remaining mtDNA impairs mitochondrial respiration, which increases oxidative stress over time, and eventually, accumulating ROS damages the nuclear genome.

## Discussion

Previous studies show that EndoG is involved in degrading mtDNA during *Drosophila* spermatogenesis. However, EndoG only nicks dsDNA at (dG)_n_/(dC)_n_ tracks, whereas metazoan mitochondrial genome possess an unusually biased A/T composition ^22 23 24 25^, suggesting that other nucleases must be involved. Here, we discover that ExoA is a mitochondrial exonuclease, highly expressed in *Drosophila* testis, and essential for mtDNA removal in elongated spermatids. ExoA degrades DNA in both directions and prefers dA/dT nucleotides. The *exoA* and *endoG* double mutant exhibited a stronger phenotype than either mutant individually, suggesting that these two enzymes may work in parallel in mtDNA removal. Alternatively, EndoG and ExoA may work in tandem, with EndoG generating initial breaks on mtDNA, followed by ExoA degrading the nicked DNA. The removal of mtDNA is delayed but eventually executed in *endoG* mutant flies, which exhibit normal fertility ^2^, suggesting that other mechanisms independent of EndoG may generate breaks on mtDNA, facilitating the degradation of mtDNA by ExoA (Fig. 5e).

Developmental declines in mtDNA content have also been observed during sperm development in mammals ^4 26 27 28^, albeit to varying extents. The expression of human ortholog of ExoA, hPoldip2 can restore the male fertility of *exoA^del^* flies, suggesting a potentially conserved function of ExoA homologs in mammals. Unlike ExoA, mammalian Poldip2 is widely expressed across various tissues. It is possible that these proteins may have evolved a broader role in mtDNA homeostasis, extending beyond their primary function in spermatogenesis. Mitochondrial genomes have a substantially higher mutation rate than nuclear genomes in metazoans ^29^. However, the repertoire of repair pathways for mtDNA is limited ^30^. Studies in mammalian cells have demonstrated that damaged mtDNA molecules are rapidly degraded ^31 32^, followed by repopulation with intact genomes. Given the presence of multiple copies of the mitochondrial genome in each cell and the relatively higher cost of repair mechanisms ^30^, degradation of damaged mtDNA might be an effective way to preserve fidelity. It would be intriguing to investigate the potential roles of Poldip2 in regulating mtDNA homeostasis and maintaining mtDNA integrity.

In multicellular organisms, the egg is furnished with maternally derived organelles and macromolecules, including RNAs and proteins, deposited with defined polarity and spatial patterns ^33^ to support the rapid early embryonic cycles and instruct the subsequent pattern formation. In contrast, the mature sperm is a “stripped down” cell. All macromolecules and organelles in the cytoplasm, including ribosomes, ER, and Golgi, except for mitochondria and axoneme, are cleared ^34 35^. This clearance process not only prevents the deposition of sperm-derived proteins and mRNAs that could disrupt early embryonic cycles and patterning, but also streamlines sperm shape for effective movement ^33 35^. It has been demonstrated that mtDNA replisomes are enriched in two membrane-spanning structures and tethered to ER-mitochondrial contacts ^36 37^, which are frequently observed in developing spermatids ^34 38^. If not degraded, mtDNA could be linked to ICs indirectly through these structures in individualizing spermatids, impeding the progression of traveling ICs. This proposition explains the association of nucleoids with ICs in the *exoA* and *endoG* double mutants and the individualization defects in these flies. Persistent mtDNA could also cause potential mito-nuclear imbalance. In mature sperm, nuclear genes’ expression is completely shut down. The persistent mtDNA could produce excessive, unassembled mtDNA-encoded electron transport chain subunits, which may impair mitochondrial respiration and lead to the generation of excessive damaging free radicals ^39 40^. Supporting this idea, mature sperms of old *exoA^del^* flies exhibited an increased ROS level, along with markedly fragmented nuclear genome. A study in mice also showed that sperm mobility is negatively correlated with mtDNA copy numbers ^26^. Therefore, mtDNA removal appears essential for two key aspects of male reproductive biology: the effective removal of cytoplasm during sperm development and preventing potential mito-nuclear imbalance in mature sperms. The stringent uniparental inheritance of mitochondrial genome, one of the most mysterious genetic phenomena in multicellular organisms, might be a prerequisite for the asymmetry between two gametes in sexual reproduction.

## Acknowledgements

We thank Dr. S. Deluca for advice on project; Dr. X. Chen for proving FRT-SV40 PolyA-FRT pBluescript KS (-) plasmid; Bloomington Drosophila Stock Center; Drosophila Genomics Resource Center for various plasmids; BestGene for Drosophila injection service. This work is supported by the National Heart, Lung, and Blood Institute Intramural Research Program.

## Author contributions

Z.C. conceived and developed the concept, tested hypothesis, designed and performed the experiments, analyzed data, interpreted results, and wrote and edited the manuscript; F. Z. designed and performed ROS stanning, and edited the manuscript; A.L. contributed to the generation and the maintenance of transgenic flies; M.Y. contributed to the generation the maintenance of fly stocks; Z.W. helped with the sperm bundle isolation and staining; G.Z. helped with the TEM experiment; C.C. advised and helped with the super resolution microscopy analysis; H.X. conceived developed the concept; formulated the hypothesis; supervised the project and experimental design, interpreted results, and wrote and edited the manuscript.

## Competing interests

Authors declare no competing interests.

## Materials & Correspondence

All original data needed to evaluate the manuscript are deposited in the NIH shared drive, and will be available upon request, following the Journal’s instruction. All materials presented in this work will be available upon request, under a material transfer agreement with NHLBI. Correspondence and material request should be addressed to H.X.

## Materials and Methods

### Fly stocks and husbandry

*Drosophila melanogaster* was reared in a humidity-controlled incubator under a 12-hour light-dark cycle at 25°C using standard cornmeal molasses agar media. The following fly stocks were used: *w^1118^*, *EndoG^MB07150^* (Bloomington Drosophila Stock Center, BDSC, stock 26072), *UAS-FLP* (BDSC, stock 62139), *Bam-gal4* ^41^, *w^1118^* (*mt:ND2^del1^*) ^42^. The *w^1118^*strain was used as control fly unless otherwise specified.

### Molecular cloning and transgenic flies

To construct ExoA-mCherry plasmid that expressing in *Drosophila* S2 cell culture, the coding sequence of *exoA* was PCR amplified from the DGRC (Drosophila Genomics Resource Center) Gold collection (DGRC, stock 3372), fused with C-terminal mCherry tags, and cloned into the pENTR3C vector (Thermo Scientific). Subsequently, the plasmids were recombined into the pAW vector (DGRC, stock 1127) using Gateway LR-clonase II (Thermo Scientific). For the ExoA-mCherry-nGFP plasmid, the *exoA* coding sequence was fused with a C-terminal mCherry tag followed by the N-terminal half of GFP sequence and inserted into the pIB-V5 vector (Thermo Scientific). Plasmid SOD2-cGFP was previously published ^43^. ExoA-6His construct was generated by cloning the *exoA* coding sequence into the pET21b vector, fused with a C-terminal 6His tag.

To generate the *exoA* mini gene, approximately 5 kb of 5’ and 3’ flanking sequences of the *exoA* gene were PCR amplified from *w^1118^*genomic DNA and cloned into the pattB vector (DGRC, stock 1420) using the in-fusion cloning method (Takara). The resulting construct was designated as pattB-exoA-mini. For the *hPoldip2* genomic rescue transgene construct, the coding region of *exoA* was substituted with that of *hPoldip2* (GeneCopoeia) on the pattB-exoA-mini plasmid. Subsequently, the rescue transgenes were produced by injecting the plasmids into 25C6 at the attP40 site via PhiC31 integrase-mediated transgenesis (BestGene).

### Generation of CRISPR knock-out and knock-in flies

The knock-out and knock-in lines, including *exoA^del^*, *endoG^KO^*, ExoA-mNeonGreen and *exoA^mitoExoIII^*, were generated using standard CRISPR methods ^44^. For the knock-out lines *exoA^del^* and *endoG^KO^*, the non-homologous end joining (NHEJ) approach was employed, involving the design of two guide RNAs (gRNAs) to delete most of the coding region of the respective genes. The guide RNA recognition sites for *exoA* deletion were GCGATGTCCCATGCTCCACAGGG (table. S2, gRNA1) and GCGCGACGATAGCGATTAAGAGG (table. S2, gRNA2), while those for *endoG* deletion were GGCAGCCGAAACAGTTCCAAAGG (table. S2, gRNA3) and GAGAGCGTGGAACGCTCGGCGGG (table. S2, gRNA4). The synthesized gRNA sequences were annealed and cloned into pU6-BbsI-chiRNA vector (Addgene #45946). Subsequently, the two plasmids carrying the gRNAs for each gene were injected into Vasa-Cas9 (BDSC, #51323) expressing embryos (BestGene). Eclosed flies were crossed with *w^1118^* flies, and the progeny carrying the deletions were screened using PCR. Sanger sequencing was performed to verify the deleted sequence.

For the ExoA-mNeonGreen knock-in line, the *mNeonGreen* coding sequence was inserted into the C-terminus of the *exoA* gene at the genomic locus via the homolog-directed repair mechanism of CRISPR. A homology donor containing a 1000-base pair upstream of the stop codon of *exoA* (left homology arm), a linker sequence ^45^, followed by the *mNeonGreen* coding sequence, and a 1000-base pair downstream of the stop codon of *exoA* (right homology arm), was cloned into the pOT2 vector. The resulting donor plasmid was co-injected with the guide RNA plasmid (table. S2, gRNA2) into Vasa-Cas9 (BDSC, #51323) expressing embryos (BestGene). Eclosed flies were crossed with *w^1118^* flies, and the progeny carrying the *mNeonGreen* insertions were screened via PCR. The sequences of the *exoA* genomic region and *mNeonGreen* were verified using Sanger sequencing.

The *exoA^mitoExoIII^* line was generated by replacing the coding region of the *exoA* gene with the FRT-SV40-FRT-mitoExoIII cassette through CRISPR. Initially, a homology donor spanning 1000-base pair upstream of the start codon (left homology arm) and the 1000-base pair downstream of the stop codon (right homology arm) of the *exoA* gene was PCR amplified and cloned into pOT2 vector. Concurrently, the FRT-SV40 PolyA-FRT plasmid/pBluescript KS (-) ^46^ was utilized, with a citrate synthase mitochondrial-targeting sequence (MTS, mito) followed by the *E.coli xthA* gene (coding Exonuclease III) cloned after the second FRT site via in-fusion cloning (Takara). Subsequently, the entire FRT-SV40-FRT-mitoExoIII sequence was recovered using NotI and EcoRI digestion, replacing the *exoA* coding region on the homology donor plasmid via in-fusion cloning. The resulting donor plasmid was then injected along with two guide RNA plasmids (table. S2, gRNA1 and gRNA2) into Vasa-Cas9 (BDSC, #51323) embryos (BestGene). Eclosed flies were crossed with *w^1118^*flies, and the progeny carrying the FRT-SV40-FRT-mitoExoIII cassette insertions were screened via PCR. The sequence of the *exoA* genomic region and the insertions were verified using Sanger sequencing.

To ensure a homogenous genetic background, all CRISPR lines were backcrossed at least six generations into a *w^1118^* background.

### Male fertility assay

Individual 2-day-old males of the specified genotypes were paired with two virgin *w^1118^* females. Every 3 days, the male flies were transferred to fresh vials and paired with another two virgin *w^1118^* females. The adult progeny from each male fly were counted. A minimum of 20 males per genotype were used for testing.

### Antibody

The antibodies used in this study were as follows: A custom antibody against ExoA was generated in rabbits using purified full-length protein as the antigen (GenScript). Mouse Polyglycylated-tubulin antibody (1:1000, MABS276, Sigma-Aldrich); Rabbit Anti-BrdU (1:200, ab152095, abcam); Rabbit Anti-hPoldip2 (1:500, 15080-1-AP, proteintech); Alexa Fluor 647 Phalloidin (1:50, A22287, Invitrogen); Alexa Fluor 568 goat α-mouse IgG (1:200, Invitrogen); and Alexa Fluor 568 goat α-rabbit IgG (1:200, Invitrogen).

### Immunostaining of *Drosophila* testes

*Drosophila* testes with the specified genotypes were dissected in Schneider’s *Drosophila* medium (21720, Gibco) supplemented with 10% fetal bovine serum (FBS) and fixed in PBS containing 4% paraformaldehyde (Electron Microscopy Sciences) for 20 minutes. After three washes with PBS, the tissues were permeabilized in 0.5%Triton X-100 in PBS for 20 minutes. Subsequently, the testes were incubated with blocking solution (PBS, 0.2% BSA, 0.1% Triton X-100) for 1 hour before being incubated with primary antibodies diluted in blocking solution at 4°C overnight. Following three washes with blocking solution, the tissues were incubated with Alexa Fluor 647 Phalloidin and Alexa Fluor-conjugated secondary antibodies for 1 hour at room temperature. Finally, the testes were mounted in Vectashield mounting medium with DAPI (H-1500, Vector Laboratories). Images were acquired using a Leica SP8 confocal system (Leica HC PL APO 63×/1.4 oil lens; LAS X acquisition software version 3.5.7; scan speed 400 Hz; Pinhole 1 A.U; Excitation 405, 488, 561 and 640 nm; z-stacks with 1 μm per step). A tile scan was performed to obtain stitched images of whole testes and seminal vesicles. Image processing was performed using Fiji software (version 2.14.0, NIH).

### Cell culture

S2 cells were cultured in Schneider’s *Drosophila* medium supplemented with 10% FBS and penicillin– streptomycin (100 U/mL) following standard procedures. One day before transfection, S2 cells were seeded onto eight-well glass bottom chambered coverslips (Lab-Tek II, Nunc) pre-treated with a 0.5 mg/ml Concanavalin A (Sigma) solution. Plasmids were transfected into S2 cells with Effectene transfection reagent (301425, Qiagen). Fluorescent images were acquired using a Perkin Elmer Ultraview confocal system (Zeiss Plan-apochromat 63×/1.4 oil lens; Volocity acquisition software; Hamamatsu Digital Camera C10600 ORCA-R2) approximately 48 hours after transfection.

To stain mitochondrial nucleoids in S2 cells, the Picogreen reagent (P11496, Invitrogen) was diluted 1:300 in the cell culture medium and incubated with the cells for 30 minutes (protected from light). After rinsing with PBS three times, fluorescent images were obtained.

### Staining of spermatid cysts and quantification of mitochondrial nucleoids

Isolation and staining of spermatid cysts were carried out following a previously published protocol with minor modifications ^2^. Testes were dissected from 2- to 3-day old male flies in ice-old TB buffer (10 mM Tris pH 6.8, 183 mM KCl, 47 mM NaCl, 1 mM EDTA) and transferred to a small drop of TB (approximately 10 μl) on a siliconized coverslip (HR3-215, Hampton Research). The base of the testis was gently torn off using forceps. While holding the unopened anterior tip of the testis with forceps, the contents were extruded using a glass capillary tube as a squeegee. The sample was then sandwiched between a poly-L-lysine (0.01%, Sigma) treated slide and the coverslip. The sandwich was briefly frozen in liquid nitrogen for 20-30 seconds, the coverslip was removed with a razor blade, and the slide containing the samples was incubated in ice-cold absolute ethanol. The tissues were fixed with 3.7% paraformaldehyde in PBS for 20 minutes, washed twice with PBS for 5 minutes each, and permeabilized with 0.1% Triton X-100 in PBS for 30 minutes. After washing in PBS twice, the samples were incubated with Alexa Fluor 647 Phalloidin diluted in blocking solution (PBS, 0.2% BSA, 0.1% Triton X-100) for 2 hours at 37°C in a humid chamber. Samples were washed three times with PBS, stained with DAPI (1 μg/ml in PBS, Sigma) for 30 minutes at room temperature, followed by washing in PBS three times and mounting in Vectashield mounting medium with DAPI (H-1500, Vector Laboratories). Images were acquired on a Perkin Elmer Ultraview confocal system (Zeiss Plan-apochromat 63×/1.4 oil lens; Volocity acquisition software; Hamamatsu Digital Camera C10600 ORCA-R2; z-stacks with 0.5 μm per step). A tile scan was performed to obtain stitched images of whole spermatid cysts.

To assess the numbers and size of mitochondrial nucleoids in spermatid bundles, image analysis was performed using Fiji (NIH). Initially, various regions of interest (ROIs) along the length of a spermatid bundle were chosen and duplicated as *z*-stack images. The length of the spermatid bundle in each ROI, as well as the distance of the ROI from the nuclear head, was measured using “measure” function. Subsequently, within each ROI, the spermatid cyst was outlined, and the “clear outside” function was applied to focus solely on the area within the spermatid cysts for analysis. Next, in the 405 nm (DAPI) channel, individual mitochondrial nucleoids were segmented using the Fiji plugin “trainable Weka segmentation 3D”. The resulting hyperstack “probability maps” were further analyzed using the “3D objects counter” function. This enabled the quantification of the numbers of segmented mitochondrial nucleoids, as well as the volume of each individual nucleoid within the ROI. The nucleoid density was calculated by dividing the total number or total volume of mitochondrial nucleoids by the length of the spermatid bundle in the respective ROI. To determine the total numbers or volumes of mitochondrial nucleoids in each spermatid bundle, nucleoid density at various points along the length of the bundle were plotted (Fig. 2e and Extended Data Fig. 2b) using GraphPad Prism (version 10.1.1). The area under the curve, which represents the total nucleoid numbers or volumes in 64 spermatids, was subsequently calculated.

### Quantification of paternal mtDNA copy number using droplet digital PCR (ddPCR)

To quantify mtDNA copy number in mature sperm, *w^1118^* (*mt:ND2^del1^*) females were crossed with *w^1118^* (*mt:wt*) or *exoA^del^* (*mt:wt*) males. For each cross, 30 2-day-old virgin females were mated with 30 2-day-old virgin males for 24 hours at 25°C. Spermatheca, the sperm storage organ, from mated or virgin control *w^1118^* (*mt:ND2^del1^*) females, was dissected in PBS. Tissues were promptly transferred to the ATL lysis buffer from QIAamp DNA micro kit (56304, Qiagen), and total DNA was extracted following the manufacturer’s instructions. To detect paternal mtDNA in embryos, 2-day old *w^1118^* (*mt:ND2^del1^*) females were crossed to 2-day-old *w^1118^* (*mt:wt*), *exoA^del^*(*mt:wt*), or control *w^1118^* (*mt:ND2^del1^*) males. Embryos were collected for 30 minutes on standard grape juice agar plates, and total DNA was extracted from the eggs using QIAamp DNA micro kit at indicated developmental time points.

Droplet digital PCR (ddPCR) was employed to quantify paternal mtDNA copy number and the number of sperm. Due to the expected much higher abundance of mtDNA compared to single-copy nuclear genes, quantification using the duplex method was deemed unreliable. Therefore, a simplex ddPCR approach was used, where mtDNA and nuDNA assays were analyzed separately using different amounts of input DNA. To specifically detect paternal wild-type mtDNA (*mt: wt*) without detecting maternal mtDNA carrying the 9-bp deletion (*mt:ND2^del1^*), a primer pair and a double quenched FAM-labelled probe were designed. Additionally, a primer pair and a 3’ quenched FAM-labelled probe targeting the Y-chromosome gene *kl-2* were designed to quantify the sperm numbers. Primers/probe sets (IdtDNA Technologies) used in the ddPCR reaction are listed in the table S3.

Total DNA from *Drosophila* tissues was digested with EcoRI enzyme at 37°C for 1 hour. Subsequently, a ddPCR reaction mix containing 1× ddPCR Supermix for Probes (186-3023, Bio-Rad), 250 nM of probe, 900 nM of each primer and the DNA template (as indicated in figure legends), was assembled. The reactions were conducted in the QX200 ddPCR system (Bio-Rad), which includes droplet generation using QX200 droplet generator, PCR reaction on a C1000 Touch thermal cycler, and analysis on a droplet reader. The cycling conditions were as follows: For mt:ND2 detection, one cycle of 95 °C for 10 minutes, 42 cycles of 95 °C (2 °C /second ramp) for 30 seconds, 51°C (2 °C /second ramp) for 1 minute, 72 °C (2°C /second ramp) for 15 seconds, one cycle of 98 °C for 10 minutes, 4°C hold. For the Y-chromosome gene, one cycle of 95 °C for 10 minutes, 40 cycles of 95 °C (2 °C /second ramp) for 30 seconds, 60°C (2 °C /second ramp) for 1 minute, one cycle of 98 °C for 10 minutes, 4°C hold. The QuantaSoft analysis software (Bio-Rad) was used to acquire and analyze data.

To evaluate the specificity of the primers/probe set for mt:ND2, a reaction containing 10 ng of total DNA from *w^1118^* (*mt:ND2^del1^*) flies mixed with varying amounts (0, 0.001, 0.005, 0.01 or 0.05 ng) of total DNA from *w^1118^* (*mt:wt*) flies was performed (Extended Data Fig. 2f, f’). The coefficient of correlation (R^2^) was calculated to be 0.9947, indicating a good correlation between the input *w^1118^* (*mt:wt*) *DNA amount and the measured copy number. It is noted that although the background signal is low, the input w^1118^* (*mt:wt*) DNA amounts lower than 0.001 ng is out of the linear range. Furthermore, the background signal arising from the presence of a large amount of *w^1118^* (*mt:ND2^del1^*) DNA (from maternal mtDNA) was considered and subtracted in the calculation.

### Purification of ExoA protein

To produce recombinant ExoA protein, the *exoA* coding sequence was cloned in frame with the C-terminus 6His-tag of pET21b vector. The plasmid was transformed, and the protein was expressed in BL21(DE3) competent cells (EC0114, Thermo Scientific). Bacteria were cultured in Luria-Bertani medium at 37°C until the optical density at the wavelength of 600 nm reaches approximately 0.6. Subsequently, 0.4 mM isopropyl-β-D-thiogalactoside (IPTG) (Sigma) was added and the bacteria were cultured for an additional 20 hours at 18°C. Cells were harvested and lysed in buffer A (50 mM sodium phosphate, pH 7.4, 0.3 M NaCl, 10 mM imidazole, 5% (vol/vol) glycerol, 10 mM β-mercaptoethanol) supplemented with 1 mg/ml lysozyme (Sigma) and EDTA-free protease inhibitor (11873580001, Roche) for 1 hour on ice, followed by sonication 5 times for 5 minutes each. The cell lysates were clarified by centrifugation at 12,000 g for 30 minutes. The supernatants were first purified using affinity chromatography on a HisTrap column (17524802, Cytiva) on an ÄKTA pure protein purification system (Cytiva). The column was washed sequentially with 40 mM and 80 mM imidazole (Sigma) in buffer A, and the bound proteins were eluted with 250 mM imidazole in buffer A. The eluted protein was dialyzed against buffer B (20 mM Tris-HCl, pH 7.5, 0.1 M NaCl, 5% (v/v) glycerol, and 1 mM DTT), and then loaded onto a 5 ml HiTrap heparin column (17040703, Cytiva) that had been equilibrated with buffer B. Following washing with buffer B, ExoA protein was eluted with a 40 ml gradient of 0.1 M to 1 M NaCl in buffer B, with ExoA eluting at salt concentrations of 0.1-0.3 M. The ExoA-containing fraction was concentrated and further purified using a Superdex 200 increase 10/300 GL size-exclusion chromatography column (28990944, Cytiva) equilibrated in buffer C (20 mM Tris-Cl, pH 7.5, 20 mM NaCl, 5% (v/v) Glycerol, 1 mM DTT and 0.1 mM EDTA). The purified proteins were stored at −80°C and the protein concentration was determined using Bradford plus protein assay reagents (23238, Thermo Scientific).

### Nuclease assay

All 6-FAM-labelled DNA oligos were synthesized using the RNase-free HPLC purification method (IdtDNA Technologies). To generate the dsDNA substrate, equal molar oligonucleotides were mixed in nuclease-free duplex buffer (IdtDNA Technologies), heated to 95°C for 5 minutes, and gradually cooled to 25°C for 45 minutes. The nuclease assay was carried out using a reaction mixture containing 10 mM Tris-HCl, pH 8.3, 2.5 mM MgCl_2_, 0.5 mM CaCl_2_, 5 mM DTT, and 100 nM of 5′- or 3’-6-FAM-labelled ssDNA or dsDNA substrate (table. S4). The reaction was initiated by adding the purified ExoA protein with the concentrations indicated in the figure legends, incubated at 37°C for the indicated time and terminated by adding Novex^TM^ TBE Urea sample buffer (LC6876, Thermo Scientific). The reaction mix was denatured for 5 minutes at 75°C and resolved on a 20% denaturing polyacrylamide gel (SequaGel, EC-833, National Diagnostics) using a Model V16 vertical electrophoresis apparatus (15 cm × 17 cm × 0.8 mm, Apogee Electrophoresis) at 300 V for 2 hours. The gels were imaged on a Typhoon biomolecular imager (GE Healthcare) using the fluorescence scanner (Cy2). Remaining substrate (%) was quantified using Fiji (NIH) and calculated by normalizing the substrate band density at each time point to the band density at time point zero.

### Transmission electron microscopy (TEM)

*Drosophila* testes were dissected in Schneider’s *Drosophila* medium supplemented with 10% FBS and immediately fixed in fixation solution (2.5% glutaraldehyde, 2% formaldehyde in 0.1 M sodium cacodylate buffer) at room temperature for 5 minutes, followed by an additional fixation on ice for 1 hour. After washing in cold cacodylate buffer, the testes were postfixed with reduced 2% Osmium tetroxide (Sigma, reduced by 1.5% potassium ferrocyanide right before use) for 1 hour on ice. After washing with water, the tissues were placed in the thiocarbohydrazide (Electron Microscopy Sciences) solution for 20 minutes at room temperature. Then, the testes were fixed in 2% Osmium tetroxide for 30 minutes at room temperature, stained *en bloc* with 1% uranyl acetate (Electron Microscopy Sciences) overnight at 4°C, and further stained with Walton’s lead aspartate solution for 30 minutes at 60°C. After dehydration with ethanol series, the samples were embedded in Epson-Aradite (Electron Microscopy Sciences). The 80 nm thin sections cut by Leica EM UC6 ultramicrotome were viewed on a Tecnai T12 (FEI, Hillsboro, OR) transmission electron microscope.

### TUNEL assay

*Drosophila* testes were dissected in Schneider’s *Drosophila* medium (10% FBS) and fixed in PBS containing 4% paraformaldehyde for 20 minutes. After washing in PBS three times, the tissues were permeabilized in 0.25%Triton X-100 in PBS for 20 minutes. Then, the testes were processed following the instructions of APO-BrdU kit (AU1001, Phoenix Flow Systems, Inc). Briefly, the samples were rinsed twice in the wash buffer before being incubated in the DNA-labeling solution containing Tdt enzyme, Br-dUTP, and Tdt reaction buffer. The reaction proceeded at 37°C for 1 hour, after which the samples were rinsed twice in rinse buffer. The samples were blocked in blocking buffer (2% BSA in PBS) for 60 minutes, then incubated with the anti-BrdU antibody (1:200, ab152095, abcam) for 2 hours at room temperature or overnight at 4°C. After three washes with PBS, the samples were incubated with Alexa Fluor 647 Phalloidin and Alexa Fluor 568 goat α-rabbit IgG (1:200, Invitrogen). Finally, the samples were mounted in Vectashield antifade mounting medium with DAPI. Images were acquired on a Perkin Elmer Ultraview confocal system.

### Single-molecule fluorescence *in situ* hybridization (smFISH) of mtDNA

The labelling of mtDNA by TFAM-mNeonGreen in testes was assessed using the single-molecule fluorescence *in situ* hybridization (smFISH) assay, following established protocols ^47^. Briefly, testes from TFAM-mNeonGreen knock-in flies were dissected and fixed in fixative buffer (100 mM sodium cacodylate, pH 7.3, 100 mM sucrose, 40 mM potassium acetate, 10 mM sodium acetate, 10 mM EGTA, 5% paraformaldehyde) for 4 minutes. Subsequently, the samples underwent sequential washing steps with 2×SSCT buffer (2×SSC with 0.1% Tween-20), 2×SSCT/20% formamide, 2xSSCT/40% formamide, and 2xSSCT/50% formamide, each for 10 minutes. To make hybridization probes, 30 pairs of 5’ labeled CAL Fluor Red 590 DNA oligonucleotide primers (see reference ^47^ for sequences) were synthesized (LGC Biosearch Technologies) and used to PCR amplify DNA fragments from *w^1118^* genomic DNA. The resulting ∼300 bp PCR products from the 30 reactions were gel purified, pooled with equal molarity, and added into the hybridization solution (2xSSC, 50% formamide, 10% dextran sulfate, 2 mg/ml BSA, 10 mM vanadyl ribonucleoside complex (Sigma)). Testes were denatured in the hybridization solution containing CAL Fluor Red 590 labeled probes (5 ng/μl) at 91°C for 2 minutes, followed by overnight hybridization at 37°C. The following day, the samples were subjected to washing steps, starting with incubation in pre-warmed 2xSSCT/50% formamide solution at 37°C, followed by room temperature incubation with 2xSSCT/40% formamide, 2xSSCT/20% formamide, and finally 2xSSCT. Last, the samples were mounted in Vectashield antifade mounting medium with DAPI, and images were acquired using a Perkin Elmer Ultraview confocal system.

### Sperm mitochondrial membrane potential staining

Seminal vesicles from male flies of the indicated genotypes were dissected in Schneider’s *Drosophila* medium (10% FBS) and transferred to a slide with PBS containing 500 nM Tetramethylrhodamine (TMRM, I34361, Invitrogen) and 500 nM MitoTracker green (M7514, Invitrogen). After covering the tissue with a coverslip, sperm were extruded from seminal vesicles by gently applying force on the coverslip. Imaging was performed immediately to minimize the effects of hypoxia. Live images were captured using a Visitech instant structured illumination microscope (iSIM, BioVision) with 488-561 Dual camera acquisition mode (Olympus UPlanApo 60x /1.3 Sil oil len; Visiview acquisition software; ORCA-Flash4.0 V2 Digital CMOS camera C11440; excitation wavelength 488 nm and 561 nm; exposure time 300 millisecond). To calculate the TMRM/MitoTracker Green ratios, image analysis was performed using Fiji (NIH). First, the 488 nm and 561 nm channels were separated in a single image. Then, regions of interest (ROI) within a sperm were selected on the TMRM channel (561 nm) using the “color threshold” function. The “restore selection” function was applied to outline the same area on the corresponding MitoTracker Green channel (488 nm). The mean intensity of the ROI and the background area in each channel were obtained through the “mean gray value” using the “measure” function. Last, the intensity of TMRM and mitoTracker Green was obtained by subtracting the background intensity from the ROI intensity, respectively. The ratiometric values were generated by normalizing the mean intensity of TMRM to that of mitoTracker Green channels.

### Detection of ROS levels

Detection of ROS levels was conducted using CellROX Deep Red according to the manufacturer’s protocol (C10422, Thermo Scientific). Briefly, the testes were incubated in Schneider’s *Drosophila* medium containing 5 μM CellROX Deep Red for 45 minutes at 25°C, followed by washes with PBS and fixation with 3.7% paraformaldehyde for 15 minutes. Samples were then mounted in Vectashield antifade mounting medium with DAPI. Images were captured using a Perkin Elmer Ultraview confocal system, and CellROX intensity was quantified using Fiji (NIH). Regions of interest (ROIs) within a seminal vesicle were selected on the CellROX Deep Red channel (640 nm). The mean intensity of CellROX signal was obtained through the “measure” function on the selected ROI with background subtracted.

### Statistical analysis

Data are shown as mean ± SD (standard deviation). GraphPad Prism software 10.0 was used to generate charts and perform statistical analysis. We used multiple unpaired t test to determine significant differences between two groups. The P value was indicated by stars: ***P < 0.0001; **P < 0.001; *P < 0.05.

---

**Supplementary Table 2.**
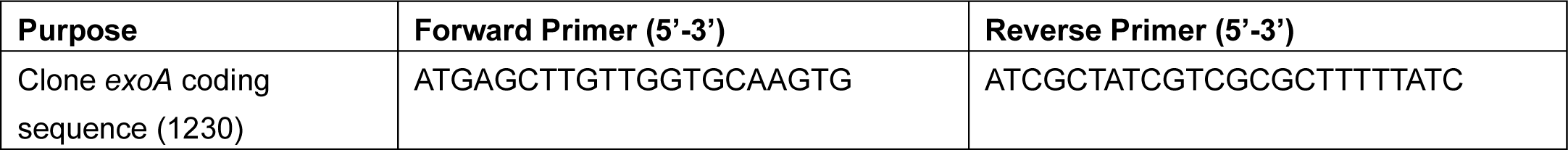
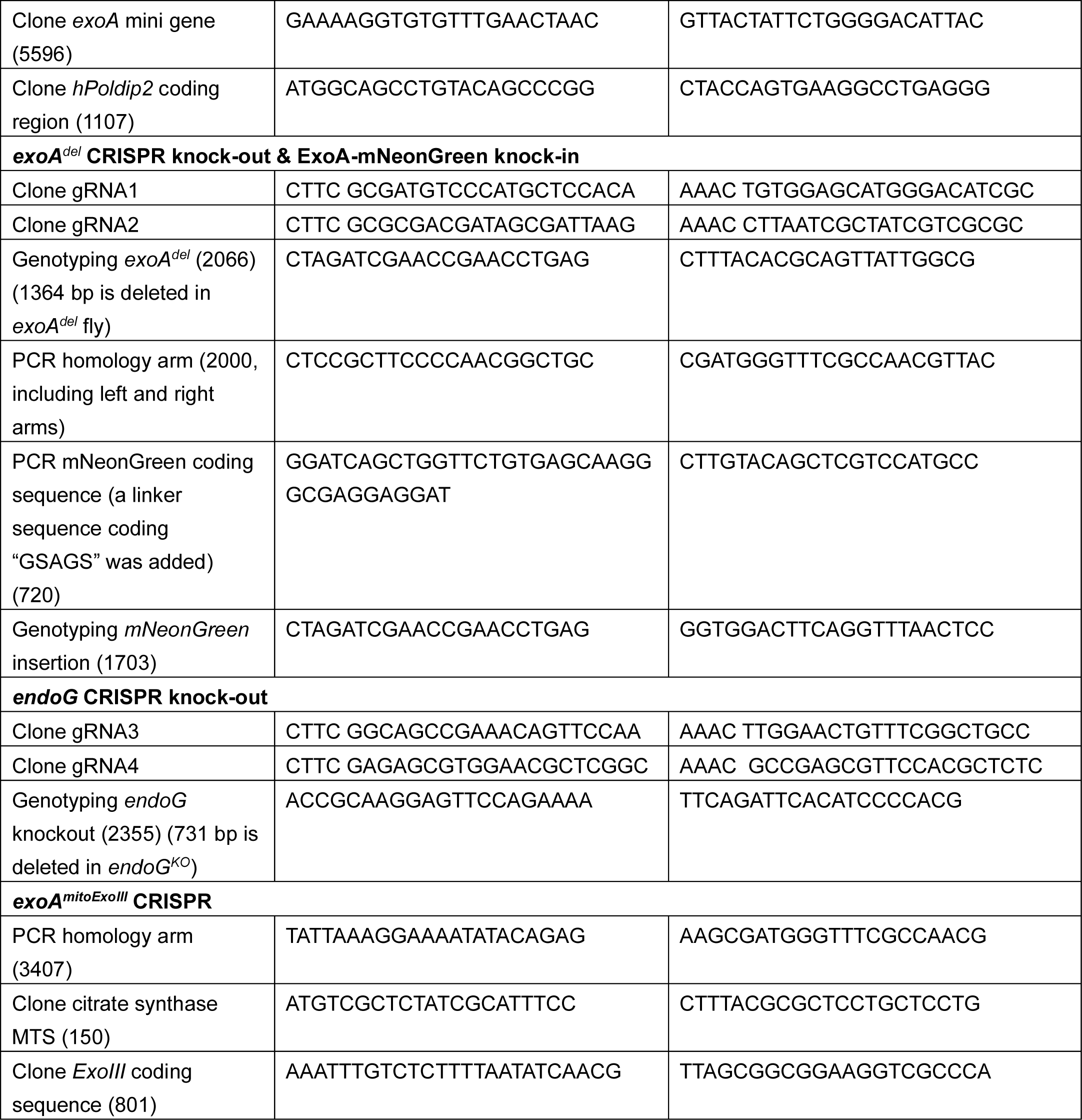
PCR primers used in this study (target length in base pairs)

**Supplementary Table 3.**
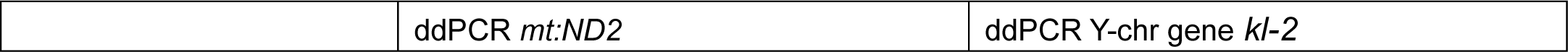
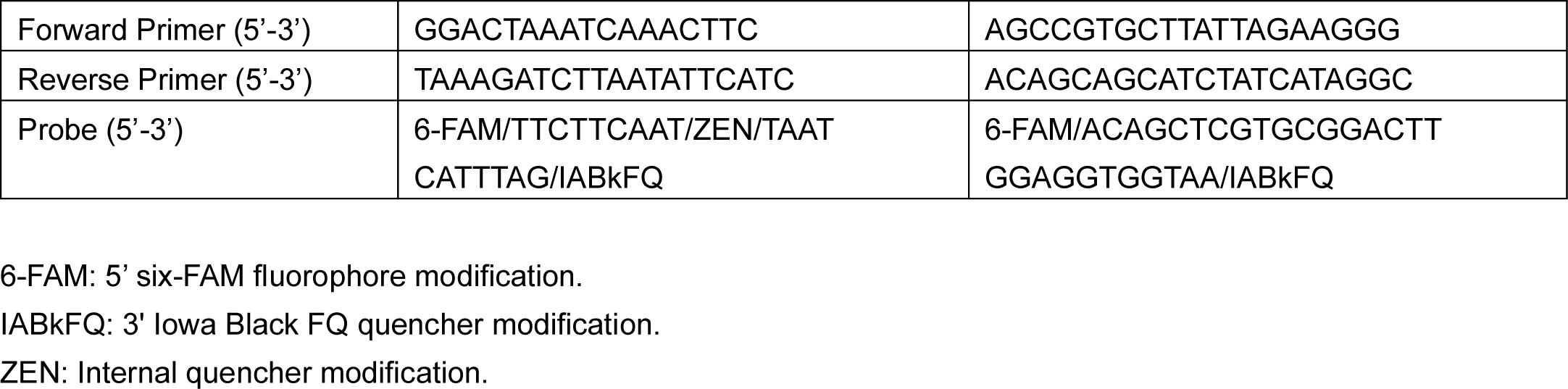
Primers/probe sets for detecting mt:ND2 and Y-chromosome gene using ddPCR.

**Supplementary Table 4.**
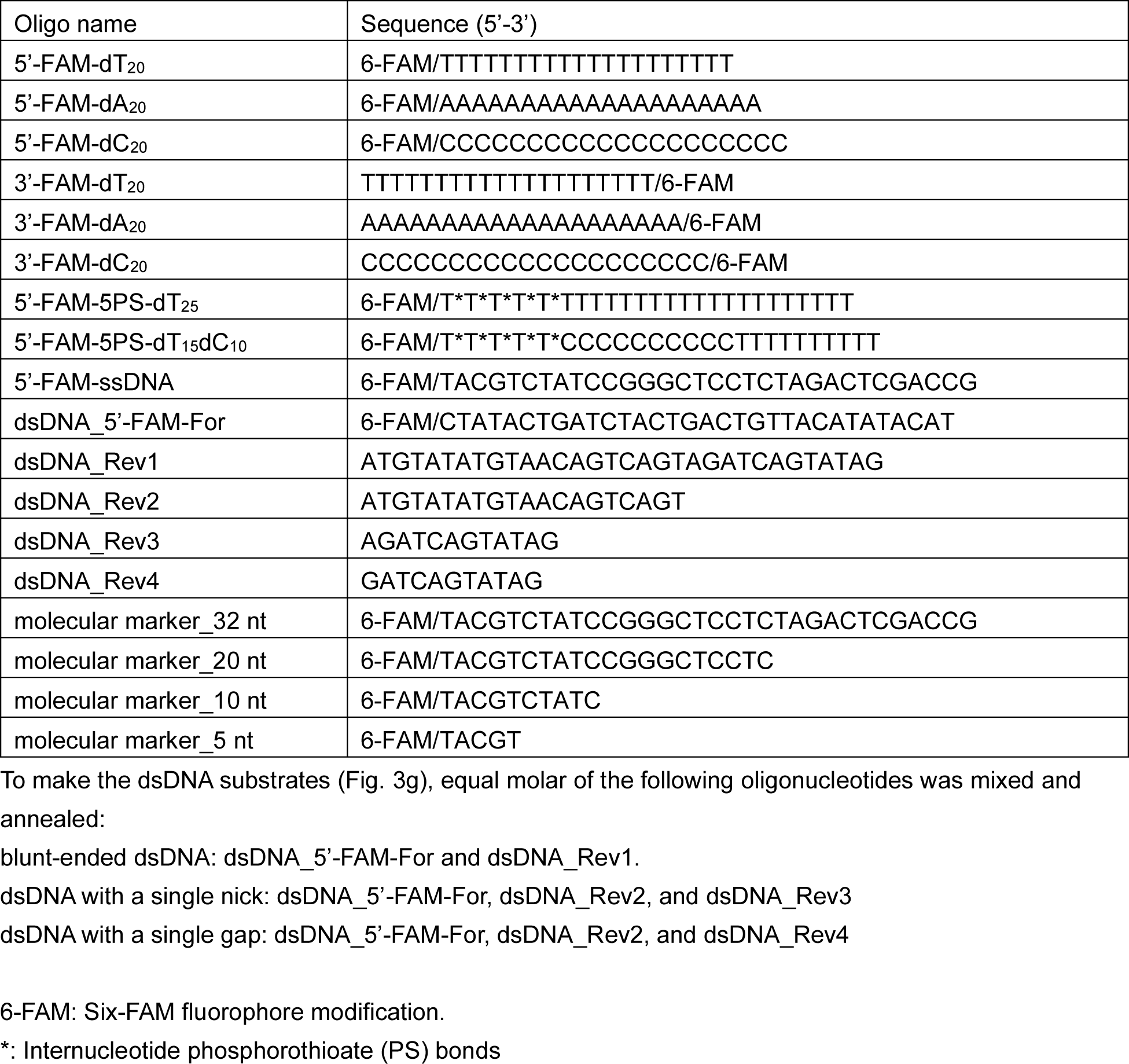
oligonucleotide sequence for nuclease assay.

## Extended Data figures and tables

**Extended Data Fig 1.**
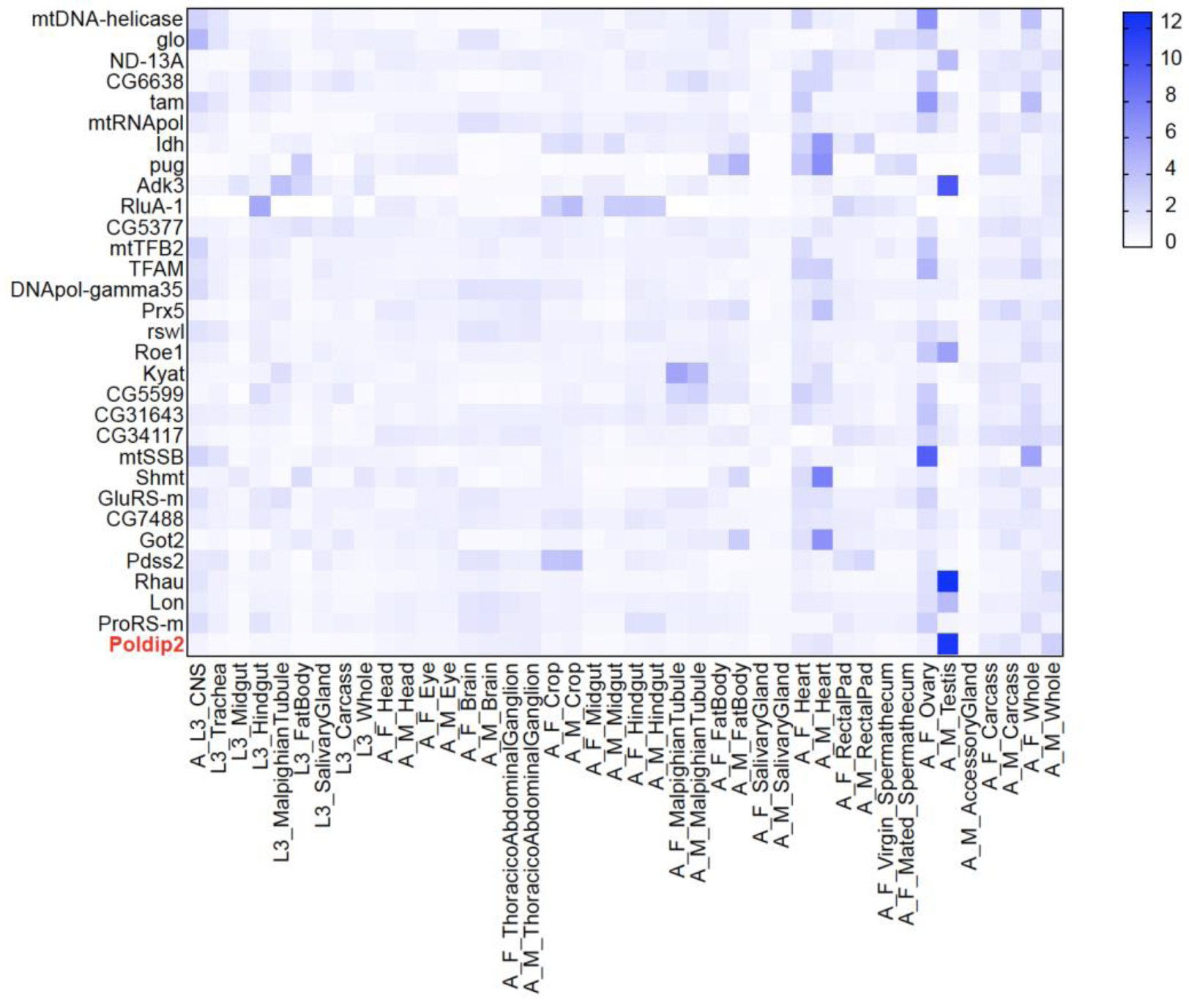
Tissue expression profile of *Drosophila* mitochondrial nucleoid associated proteins. Heatmap was generated with RNA-seq data from FlyAtlas2 (the *Drosophila* gene expression atlas). Color codes indicate the scaled RPKM (reads per kilobase per million mapped reads) folds over tissues. Note the mRNA level of ExoA/Poldip2 is significantly higher in *Drosophila* testis compared to other tissues. A, adult; L3, third instar larva; F, female; M, male.

**Extended Data Fig 2.**
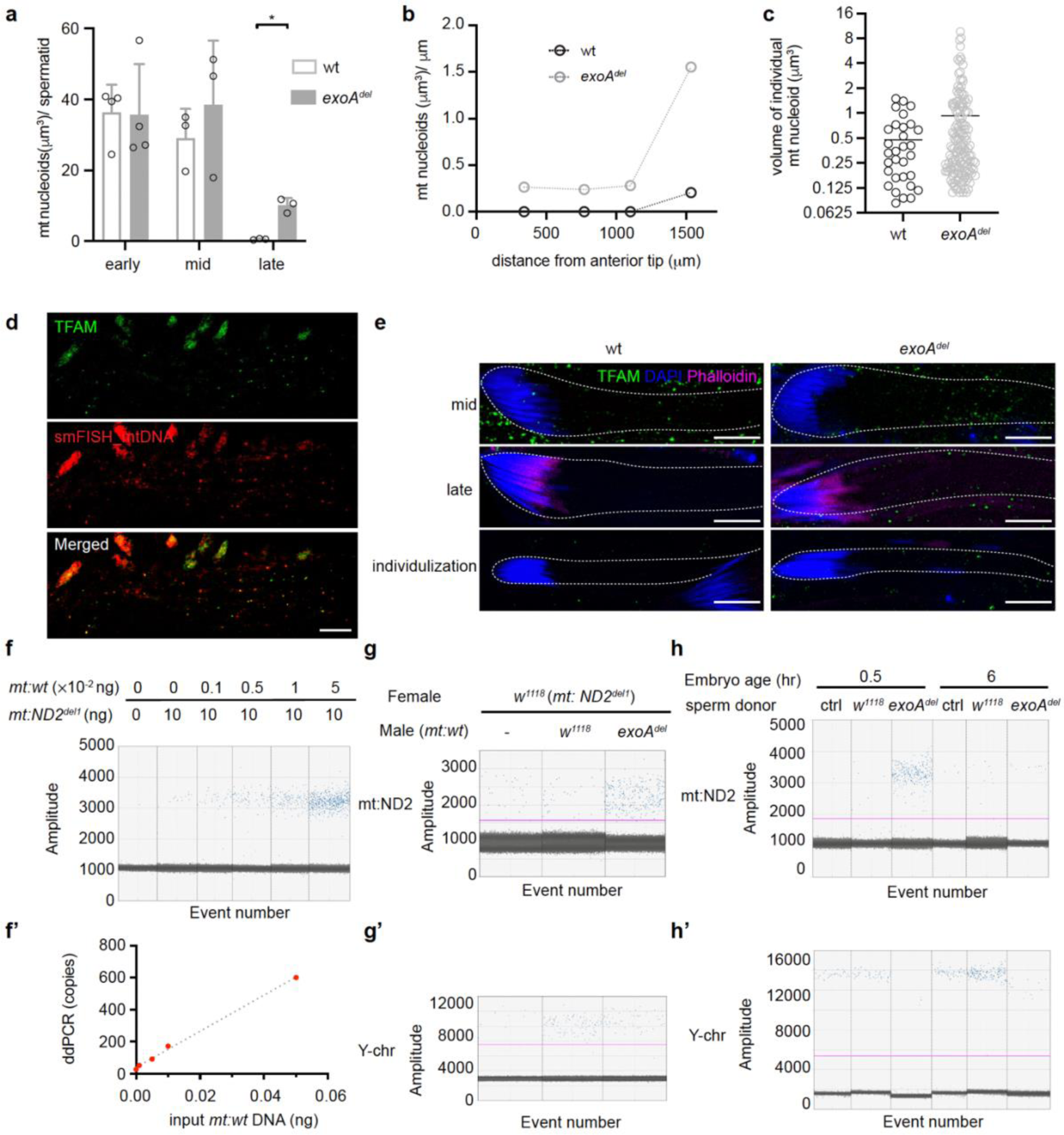
mtDNA persist in late spermatogenesis stages and mature sperm of *exoA^del^* flies. (a) Total mitochondrial nucleoids measured in volumes per spermatid at early-elongating (early), mid-elongating (mid) and fully elongated (late) stages. Error bars represent standard deviation. * P<0.05. (b) Density of mitochondrial nucleoids (total volumes per μm) along the length of a representative fully elongated spermatid bundle for *w^1118^* (wt) and *exoA^del^* flies, respectively. (c) A scatter dot plot displaying the distribution of individual mitochondrial nucleoid volumes from the elongated stage spermatids of *w^1118^* (wt) and *exoA^del^* flies. The solid lines indicate the mean volume. (d) TFAM-mNeonGreen can be used as a mitochondrial nucleoid marker in *Drosophila* testis. Single-molecule fluorescent *in situ* hybridization (smFISH) analysis using fluorescently labeled DNA probes specific for mtDNA (red) is colocalized with TFAM-mNeonGreen in the *Drosophila* testis. Scale bar, 10 μm. (e) Mitochondrial nucleoids labeled by TFAM-mNeonGreen demonstrates a consistent pattern of mtDNA elimination during spermatogenesis in both *w^1118^* (wt) and *exoA^del^* flies, compared with DNA dye staining. In elongating spermatids of both wt and *exoA^del^* testis, intense TFAM-mNeonGreen puncta signals were detected. The signals were rare in fully elongated and individualization stage spermatids of wt flies. Conversely, persistent mtDNA was frequently observed in the same stages of *exoA^del^* spermatids. Phalloidin (magenta) stains actin; DAPI (blue) stains nuDNA. Scale bar, 10 μm. (f) Evaluating the specificity of the primers/probe sets targeting mtDNA-encoded ND2 locus (*mt:ND2*) using droplet digital PCR (ddPCR) assay. A reaction containing 10 ng of total DNA from *w^1118^* (*mt:ND2^del1^*), a fly strain carrying a 9 base pair deletion on mtDNA-encoded ND2 locus, mixed with 0, 0.001, 0.005, 0.01 or 0.05 ng of total DNA from *w^1118^* (*mt:wt*, wild-type mtDNA) flies was performed. The ddPCR primers/probe were designed to target the *mt:wt*, while excluding *mt:ND2^del1^* mtDNA. (f’) Correlation of the amount of input *w^1118^* (*mt:wt*) DNA with the resulting mtDNA copy numbers using ddPCR. Simple linear regression was carried out and the coefficient of correlation R^2^ = 0.9947. (g-g’) Quantification of the mtDNA copy numbers per sperm in *w^1118^*and *exoA^del^* flies using ddPCR. Crosses were conducted between female *w^1118^* (*mt:ND2^del1^*) and male *w^1118^*(*mt:wt*) or *exoA^del^* (*mt:wt*) flies. Then the total DNA from the female spermatheca were extracted and analyzed. The virgin female *w^1118^* (*mt:ND2^del1^*) flies were used as the negative control. The primers/probe sets were designed to target *mt:ND2* (g) and Y-chromosome gene *kl-2* (g’), respectively. The input total DNA for detecting *mt:ND2* gene is 5 ng for each reaction. The input total DNA for detecting Y-chromosome gene is 125 ng for each reaction. (h-h’) Sperm-derived mtDNA can be transferred to embryos but gets eliminated quickly. Crosses were conducted between female *w^1118^* (*mt:ND2^del1^*) and male *w^1118^* (*mt:wt*) or *exoA^del^*(*mt:wt*) flies. The group of *w^1118^* (*mt:ND2^del1^*) being used as the sperm donor were negative control (ctrl). Embryos were collected within 0-30 min after being laid and were analyzed immediately (0.5 hr) or after 6 hours (6 hr) of development. The primers/probe sets were designed to target *mt:ND2* (h) and Y-chromosome gene *kl-2* (h’), respectively. The input total DNA for detecting *mt:ND2* gene is 10 ng for each reaction. The input total DNA for detecting Y-chromosome gene is 10 ng, 10 ng, 250 ng, 10 ng, 10 ng and 50 ng, respectively.

**Extended Data Fig 3.**
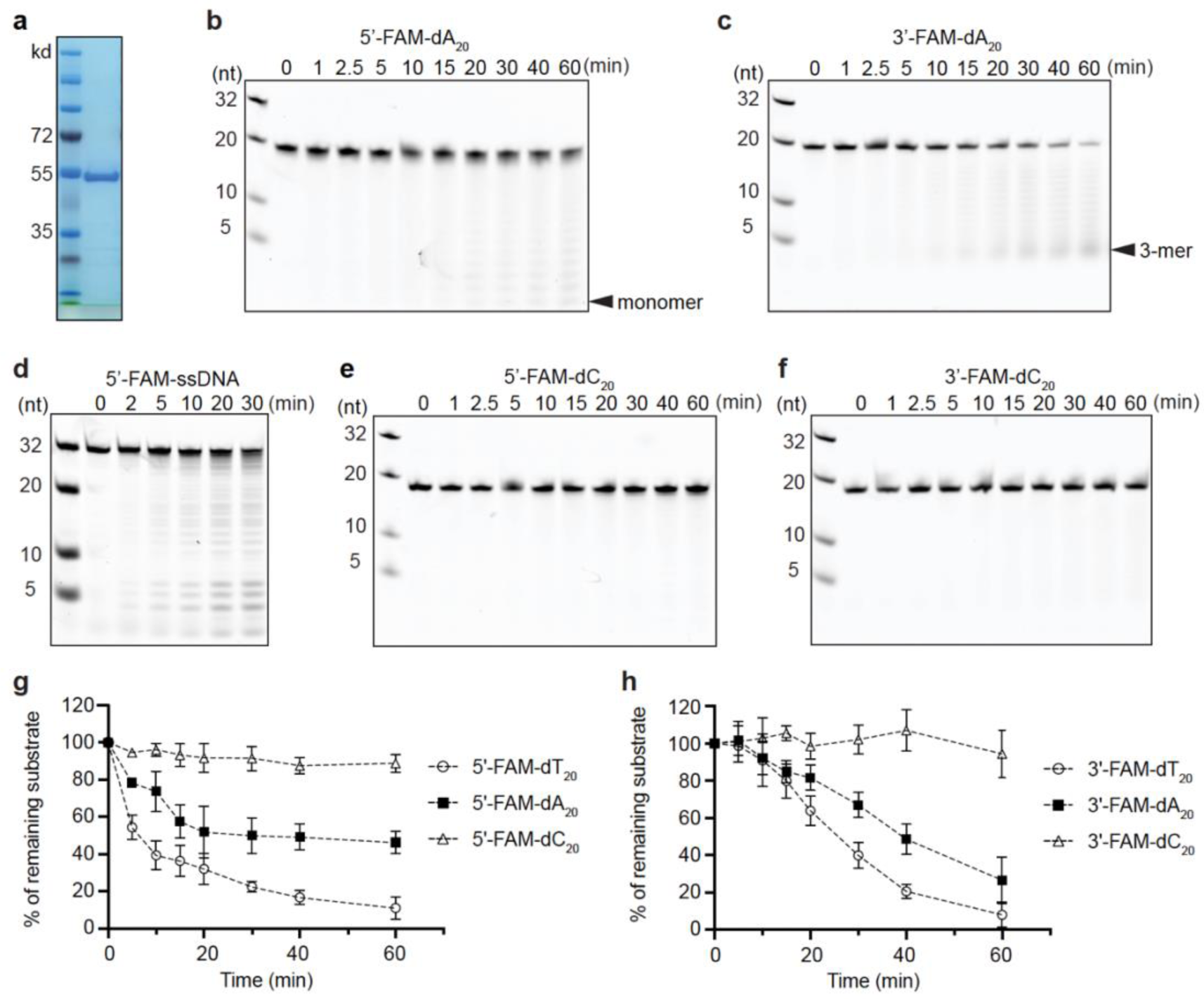
ExoA is a mitochondrial DNA exonuclease. (a) SDS-PAGE of the purified ExoA protein. M, molecular weight marker. (b, c) Degradation pattern of 5’-6-FAM and 3’-6-FAM labeled 20-nt poly(dA) single-stranded (ssDNA) substrates. The 100 nM 5’-6-FAM (b) or 3’-6-FAM (c) labeled 20-nt poly(dA) were incubated with ExoA protein (200 nM) at 37 °C and analyzed at the indicated time points. (d) Degradation pattern of a 5’-6-FAM labeled ssDNA substrate consisted of mixed dA, dT, dC and dG. The 100 nM 5’-6-FAM labeled 32-nt ssDNA was incubated with ExoA protein (200 nM) at 37 °C and analyzed at the indicated time points. (e, f) Degradation pattern of 5’-6-FAM and 3’-6-FAM labeled 20-nt poly(dC) ssDNA substrates. The 100 nM 5’-6-FAM (e) or 3’-6-FAM (f) labeled 20-nt poly(dC) were incubated with ExoA protein (200 nM) at 37 °C and analyzed at the indicated time points. (g, h) Quantification of the remaining full-length substrates, including 5’-6-FAM (g) or 3’-6-FAM (h) labeled 20-nt poly(dT), poly(dA) and poly (dC), at each time point. Data are normalized to the initial level of the full-length substrates and plotted (n=3). The molecular markers in this figure are an equal molar mixture of 5’-6-FAM labeled 32-nt, 20-nt, 10-nt and 5-nt oligonucleotides and were loaded at a concentration of 50 nM for each. Error bars represent standard deviation.

**Extended Data Fig 4.**
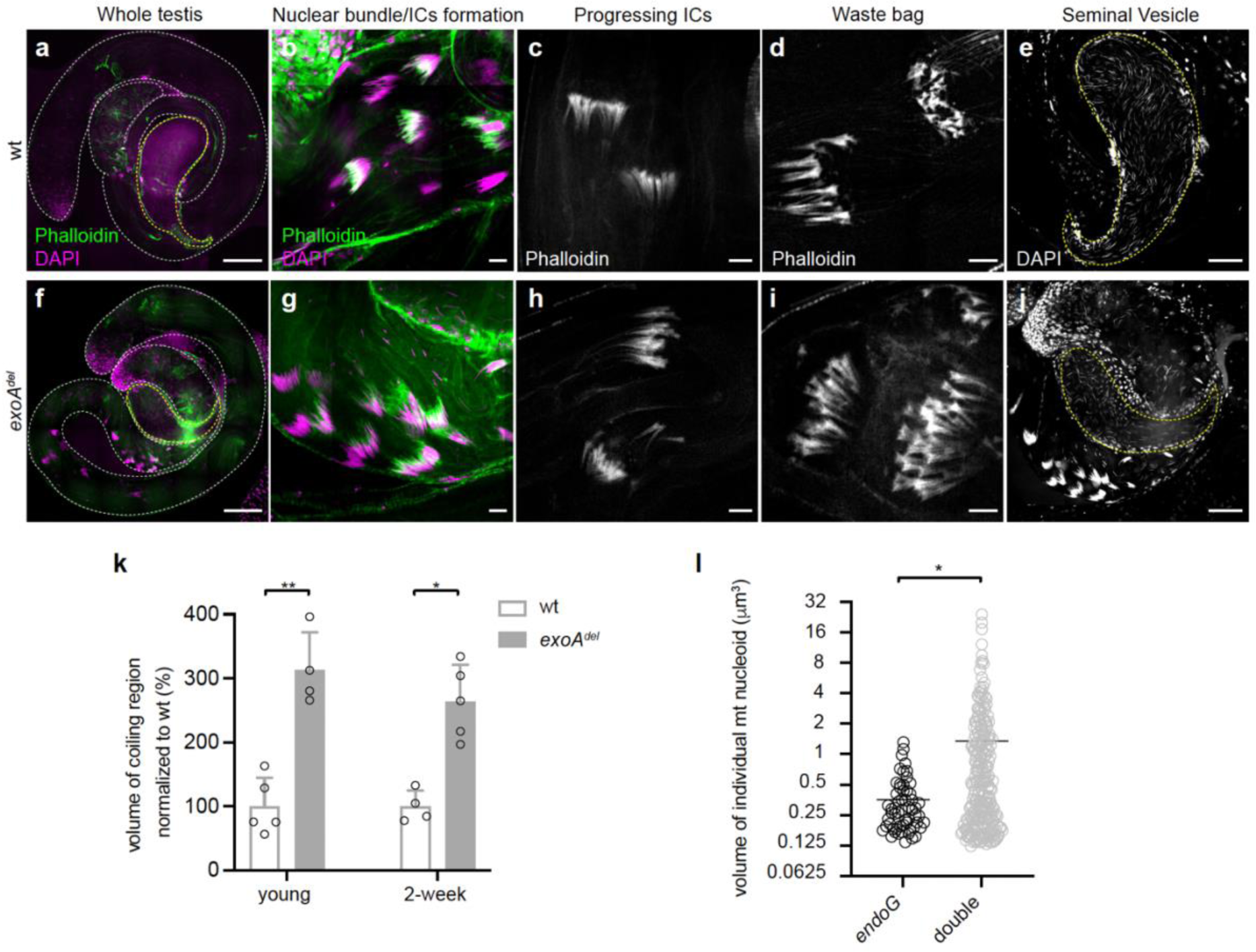
Persistent mtDNA impedes spermatid individualization. (a-j) Representative images showing the whole testis (a, f), individualization complexes (ICs) formation next to the nuclear head in the testis basal region (b, g), actin cone structures in progressing ICs (c, h), waste bags (d, i), and seminal vesicles (e, j, yellow dashed line) in *w^1118^*(wt) and *exoA^del^* flies. Phalloidin stains actin; DAPI stains nuDNA. Scale bar, 100 μm in **a** and **f**; 10 μm in **b**-**d** and **g**-**i**; 50 μm in **e** and **j**. (k) The coiling region in both young and 2-week-old *exoA^del^*flies is enlarged compared with wt control. Error bars represent standard deviation. ** P<0.001; * P<0.05. (l) A scatter dot plot displaying the distribution of individual mitochondrial nucleoid volumes from the elongated stage spermatids of *endoG* (*endoG^MB07150/KO^*) and double mutants (*endoG^MB07150/KO^*; *exoA^del^*). The solid lines indicate the mean volume. * P<0.05.

**Extended Data Fig 5.**
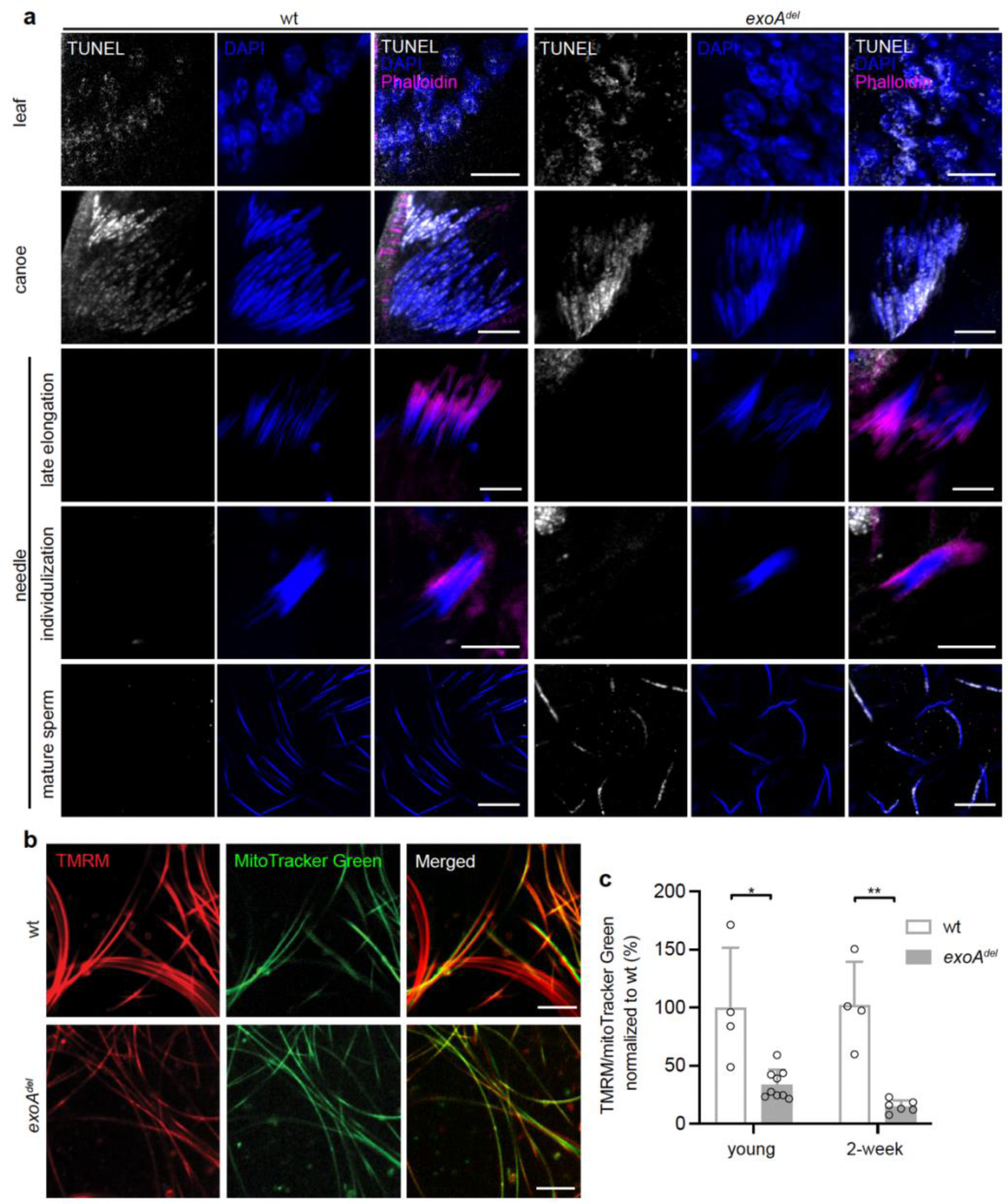
Persistent mtDNA in mature sperm causes nuclear DNA fragmentation. (a) Representative images showing nuDNA breaks labelled by TUNEL assay in the process of chromatin remodeling during spermatogenesis. The highest abundance of TUNEL signal (white) was observed in the late canoe stage, which corresponds to the histone-to-protamine transition phase. The nuDNA breaks subsequently disappeared in needle-shaped nuclei during late elongation and individualization stages, indicating repair of nuDNA breaks after the transition. No significant differences were observed between wt and *exo^del^* flies throughout this process. The developmental stages were distinguished by the morphology of nuclear heads stained with DAPI (blue), and the positioning of actin cones stained with phalloidin (magenta). Scale bar, 10 μm. (b) Compromised mitochondrial membrane potential of *exoA^del^* sperms. Mature sperm from *w^1118^* (wt) and *exoA^del^* seminal vesicles were stained with TMRM (red), a dye sensitive to mitochondrial membrane potential, in combination with mitoTracker Green (green) as a reference. Scale bar, 10 μm. (c) Quantification of TMRM/mitoTracker green ratios in both young and 2-week-old flies. Error bars represent standard deviation. ** P<0.001; * P<0.05.

**Supplementary Table 1. *Drosophila* ortholog of human mitochondrial nucleoid proteins and their tissue expression profile from FlyAtlas2 and modENCODE.**

## Notes

### Competing Interest Statement

The authors have declared no competing interest.

## References

1 Nishimura, Y. et al. Active digestion of sperm mitochondrial DNA in single living sperm revealed by optical tweezers. P Natl Acad Sci USA 103, 1382–1387 (2006). 10.1073/pnas.0506911103

2 DeLuca, S. Z. & O’Farrell, P. H. Barriers to Male Transmission of Mitochondrial DNA in Sperm Development. Developmental Cell 22, 660–668 (2012). 10.1016/j.devcel.2011.12.021

3 Yu, Z. S., O’Farrell, P. H., Yakubovich, N. & DeLuca, S. Z. The Mitochondrial DNA Polymerase Promotes Elimination of Paternal Mitochondrial Genomes. Current Biology 27, 1033–1039 (2017). 10.1016/j.cub.2017.02.014

4 Lee, W. et al. Molecular basis for maternal inheritance of human mitochondrial DNA. Nat Genet 55, 1632–1639 (2023). 10.1038/s41588-023-01505-9

5 Sutovsky, P. et al. Development - Ubiquitin tag for sperm mitochondria. Nature 402, 371–372 (1999). Doi 10.1038/46466

6 Sutovsky, P. et al. Ubiquitinated sperm mitochondria, selective proteolysis, and the regulation of mitochondrial inheritance in mammalian embryos. Biol Reprod 63, 582–590 (2000). DOI 10.1095/biolreprod63.2.582

7 Zhou, Q. H. et al. Mitochondrial endonuclease G mediates breakdown of paternal mitochondria upon fertilization. Science 353, 394–399 (2016). 10.1126/science.aaf4777

8 Politi, Y. et al. Paternal Mitochondrial Destruction after Fertilization Is Mediated by a Common Endocytic and Autophagic Pathway in Drosophila. Developmental Cell 29, 305–320 (2014). 10.1016/j.devcel.2014.04.005

9 Sato, M. & Sato, K. Degradation of Paternal Mitochondria by Fertilization-Triggered Autophagy in C. elegans Embryos. Science 334, 1141–1144 (2011). 10.1126/science.1210333

10 Gyllensten, U., Wharton, D., Josefsson, A. & Wilson, A. C. Paternal Inheritance of Mitochondrial-DNA in Mice. Nature 352, 255–257 (1991). DOI 10.1038/352255a0

11 Wolff, J. N. & Gemmell, N. J. Lost in the zygote: the dilution of paternal mtDNA upon fertilization. Heredity 101, 429–434 (2008). 10.1038/hdy.2008.74

12 Cote, J. & Ruizcarrillo, A. Primers for Mitochondrial-DNA Replication Generated by Endonuclease-G. Science 261, 765–769 (1993). DOI 10.1126/science.7688144

13 Ruizcarrillo, A. & Renaud, J. Endonuclease-G - a (Dg)N.(Dc)N-Specific Dnase from Higher Eukaryotes. Embo J 6, 401–407 (1987). DOI 10.1002/j.1460-2075.1987.tb04769.x

14 Li, L. Y., Luo, L. & Wang, X. D. Endonuclease G is an apoptotic DNase when released from mitochondria. Nature 412, 95–99 (2001). Doi 10.1038/35083620

15 Han, S. et al. Proximity Biotinylation as a Method for Mapping Proteins Associated with mtDNA in Living Cells. Cell Chem Biol 24, 404–414 (2017). 10.1016/j.chembiol.2017.02.002

16 Bogenhagen, D. F., Rousseau, D. & Burke, S. The layered structure of human mitochondrial DNA nucleoids. J Biol Chem 283, 3665–3675 (2008). 10.1074/jbc.M708444200

17 Rajala, N. et al. Whole Cell Formaldehyde Cross-Linking Simplifies Purification of Mitochondrial Nucleoids and Associated Proteins Involved in Mitochondrial Gene Expression. Plos One 10 (2015). ARTN e0116726 10.1371/journal.pone.0116726

18 Liu, L., Rodriguez-Bèlmonte, E. M., Mazloum, N., Xie, B. & Lee, M. Y. W. T. Identification of a novel protein, PDIP38, that interacts with the p50 subunit of DNA polymerase δ and proliferating cell nuclear antigen. Journal of Biological Chemistry 278, 10041–10047 (2003). 10.1074/jbc.M208694200

19 Maga, G. et al. DNA polymerase delta-interacting protein 2 is a processivity factor for DNA polymerase lambda during 8-oxo-7,8-dihydroguanine bypass. P Natl Acad Sci USA 110, 18850–18855 (2013). 10.1073/pnas.1308760110

20 Zhang, F. et al. Genetic and bioinformatic analyses reveal transcriptional networks underlying dual genomic coordination of mitochondrial biogenesis. bioRxiv (2024). 10.1101/2024.01.25.577217

21 Connelly, S. & Manley, J. L. A functional mRNA polyadenylation signal is required for transcription termination by RNA polymerase II. Genes Dev 2, 440–452 (1988). 10.1101/gad.2.4.440

22 Gammage, P. A. & Frezza, C. Mitochondrial DNA: the overlooked oncogenome? Bmc Biology 17 (2019). ARTN 53 10.1186/s12915-019-0668-y

23 Kurabayashi, A. & Ueshima, R. Complete sequence of the mitochondrial DNA of the primitive opisthobranch gastropod Pupa strigosa: Systematic implication of the genome organization. Molecular Biology and Evolution 17, 266–277 (2000). DOI 10.1093/oxfordjournals.molbev.a026306

24 Wolstenholme, D. R. Animal mitochondrial DNA: structure and evolution. Int Rev Cytol 141, 173-216 (1992). 10.1016/s0074-7696(08)62066-5

25 Chen, Z., Zhang, F. & Xu, H. Human mitochondrial DNA diseases and Drosophila models. J Genet Genomics 46, 201–212 (2019). 10.1016/j.jgg.2019.03.009

26 Luo, S. M. et al. Unique insights into maternal mitochondrial inheritance in mice. P Natl Acad Sci USA 110, 13038–13043 (2013). 10.1073/pnas.1303231110

27 May-Panloup, P. et al. Increased sperm mitochondrial DNA content in male infertility. Human Reproduction 18, 550–556 (2003). 10.1093/humrep/deg096

28 Larsson, N. G., Oldfors, A., Garman, J. D., Barsh, G. S. & Clayton, D. A. Down-regulation of mitochondrial transcription factor A during spermatogenesis in humans. Human Molecular Genetics 6, 185–191 (1997). DOI 10.1093/hmg/6.2.185

29 Lynch, M., Koskella, B. & Schaack, S. Mutation pressure and the evolution of organelle genomic architecture. Science 311, 1727–1730 (2006). 10.1126/science.1118884

30 Fu, Y., Tigano, M. & Sfeir, A. Safeguarding mitochondrial genomes in higher eukaryotes. Nature Structural & Molecular Biology 27, 687–695 (2020). 10.1038/s41594-020-0474-9

31 Srivastava, S. & Moraes, C. T. Manipulating mitochondrial DNA heteroplasmy by a mitochondrially targeted restriction endonuclease. Human Molecular Genetics 10, 3093–3099 (2001). DOI 10.1093/hmg/10.26.3093

32 Nissanka, N., Bacman, S. R., Plastini, M. J. & Moraes, C. T. The mitochondrial DNA polymerase gamma degrades linear DNA fragments precluding the formation of deletions. Nature Communications 9 (2018). ARTN 2491 10.1038/s41467-018-04895-1

33 Kimelman, D. & Martin, B. L. Anterior-posterior patterning in early development: three strategies. Wiley Interdiscip Rev Dev Biol 1, 253–266 (2012). 10.1002/wdev.25

34 Fabian, L. & Brill, J. A. Drosophila spermiogenesis: Big things come from little packages. Spermatogenesis 2, 197–212 (2012). 10.4161/spmg.21798

35 Barratt, C. L., Kay, V. & Oxenham, S. K. The human spermatozoon - a stripped down but refined machine. J Biol 8, 63 (2009). 10.1186/jbiol167

36 Meeusen, S. & Nunnari, J. Evidence for a two membrane-spanning autonomous mitochondrial DNA replisome. J Cell Biol 163, 503–510 (2003). 10.1083/jcb.200304040

37 Lewis, S. C., Uchiyama, L. F. & Nunnari, J. ER-mitochondria contacts couple mtDNA synthesis with mitochondrial division in human cells. Science 353 (2016). ARTN aaf5549 10.1126/science.aaf5549

38 Tokuyasu, K. T. Dynamics of Spermiogenesis in Drosophila-Melanogaster .3. Relation between Axoneme and Mitochondrial Derivatives. Experimental Cell Research 84, 239–250 (1974). Doi 10.1016/0014-4827(74)90402-9

39 Zhao, Q. et al. A mitochondrial specific stress response in mammalian cells. Embo J 21, 4411–4419 (2002). 10.1093/emboj/cdf445

40 Sutandy, F. X. R., Gossner, I., Tascher, G. & Munch, C. A cytosolic surveillance mechanism activates the mitochondrial UPR. Nature 618, 849–854 (2023). 10.1038/s41586-023-06142-0

41 Chen, D. & McKearin, D. M. A discrete transcriptional silencer in the bam gene determines asymmetric division of the Drosophila germline stem cell. Development 130, 1159–1170 (2003). 10.1242/dev.00325

42 Xu, H. Manipulating the metazoan mitochondrial genome with targeted restriction enzymes (vol 321, pg 575, 2008). Science 322, 1466–1466 (2008).

43 Zhang, F. et al. The cAMP phosphodiesterase Prune localizes to the mitochondrial matrix and promotes mtDNA replication by stabilizing TFAM. Embo Reports 16, 520–527 (2015). DOI 10.15252/embr.201439636

44 Gratz, S. J., Rubinstein, C. D., Harrison, M. M., Wildonger, J. & O’Connor-Giles, K. M. CRISPR-Cas9 Genome Editing in Drosophila. Curr Protoc Mol Biol 111, 31 32 31–31 32 20 (2015). 10.1002/0471142727.mb3102s111

45 Waldo, G. S., Standish, B. M., Berendzen, J. & Terwilliger, T. C. Rapid protein-folding assay using green fluorescent protein. Nat Biotechnol 17, 691–695 (1999). 10.1038/10904

46 Tran, V., Lim, C., Xie, J. & Chen, X. Asymmetric Division of Drosophila Male Germline Stem Cell Shows Asymmetric Histone Distribution. Science 338, 679–682 (2012). 10.1126/science.1226028

47 Hurd, T. R. et al. Long Oskar Controls Mitochondrial Inheritance in Drosophila melanogaster. Dev Cell 39, 560–571 (2016). 10.1016/j.devcel.2016.11.004

